# Intestinal cDC1s provide IL-12 dependent and independent functions required for CD4^+^ T cell-mediated resistance to *Cryptosporidium*

**DOI:** 10.1101/2023.11.11.566669

**Authors:** Ian S. Cohn, Bethan A. Wallbank, Breanne E. Haskins, Keenan M. O’Dea, Ryan D. Pardy, Sebastian Shaw, Maria I. Merolle, Jodi A. Gullicksrud, David A. Christian, Boris Striepen, Christopher A. Hunter

## Abstract

*Cryptosporidium* is an enteric pathogen that is a prominent cause of diarrheal disease. Control of this infection requires CD4^+^ T cells, though the processes that lead to T cell-mediated resistance have been difficult to assess. Here, *Cryptosporidium* parasites that express MHCII-restricted model antigens were generated to dissect the early events that influence CD4^+^ T cell priming and effector function. These studies highlight that parasite-specific CD4^+^ T cells are primed in the draining mesenteric lymph node (mesLN) and differentiate into Th1 cells in the gut, where they mediate IFN-γ-dependent control of the infection. Although type 1 conventional dendritic cells (cDC1s) were not required for initial priming of CD4^+^ T cells, cDC1s were required for CD4^+^ T cell expansion and gut homing. cDC1s were also a major source of IL-12 that was not required for priming but promoted full differentiation of CD4^+^ T cells and local production of IFN-γ. Together, these studies reveal distinct roles for cDC1s in shaping CD4^+^ T cell responses to enteric infection: first to drive early expansion in the mesLN and second to drive effector responses in the gut.

**Summary:** *Cryptosporidium* parasites that express model antigens were generated to dissect how parasite-specific CD4^+^ T cells are primed and mediate effector functions required to control this enteric pathogen. cDC1s produced IL-12p40 and were required for early expansion and gut homing of CD4^+^ T cells. However, IL-12p40 was only required for the development of Th1 CD4^+^ T cell effector function in the gut.

## Introduction

Conventional dendritic cells (cDCs) are professional antigen presenting cells (APCs) located in peripheral and lymphatic tissues that serve as an intermediary between the innate and adaptive immune systems. Relevant DC functions include the ability to detect microbial ligands and danger signals, which can lead to cDC production of cytokines, up-regulation of costimulatory molecules, and MHC-restricted presentation of peptide antigens to CD4^+^ and CD8^+^ T cells (1). cDCs can be defined based on developmental origin as either type 1 (cDC1) or type 2 (cDC2), where cDC1s rely on the transcription factors BATF3 and IRF8 for development and are specialized to cross-present antigens to CD8^+^ T cells. In contrast cDC2s rely on the transcription factor IRF4 and prime CD4^+^ T cells (1). This division of labor is based in part on early studies where cDC2s were shown to be superior at MHCII presentation of soluble ovalbumin (OVA) or OVA complexed to antibodies targeting DC surface receptors (2,3). However, there are examples that do not easily fit this paradigm that include the presentation of cell-associated antigens derived from tumors or intracellular infections that drive Th1 responses (4). Thus, studies have reported poorer MHCII antigen presentation by cDC2s when compared to cDC1s in these contexts (5,6). Indeed, cDC1s express MHCII and are also known to produce cytokines such as IL-12 and IL-27 that can promote Th1 CD4^+^ T cell responses (7,8). The rules that govern this division of labor and how they relate to different infections with tropisms for diverse host cell populations is not well defined.

In the gut, cDC1s and cDC2s have been shown to have distinct roles in the regulation of inflammatory and tolerogenic CD4^+^ responses. For example, cDC1s can cross-present epithelial-derived antigens to CD8^+^ T cells, provide signals necessary for maintenance of intraepithelial lymphocytes (IELs), and maintain homeostatic Th1 CD4^+^ T cells (8–10). Conversely, cDC2s are necessary for induction of anti-helminth Th2 responses (11) and Th17 responses during homeostasis and against fungal pathogens (12,13). Additionally, unique populations of APCs exist in the gut associated with the development of tolerance (14), whose role in infection-induced CD4^+^ responses remains poorly understood. The presence of these diverse APC subsets and their distinct functions likely reflects the need for the intestine to regulate CD4^+^ T cell functions to respond to diverse microbial challenges while being able to maintain tolerance. It is therefore critical to understand how responses to enteric pathogens are regulated in order to respond to the infectious challenge appropriately.

*Cryptosporidium* species are important causes of diarrheal illness globally and are associated with significant morbidity and mortality (15,16). Fecal-oral transmission of these parasites causes self-limited disease in immunocompetent individuals, which can be life-threatening in the immunocompromised host (17–21). *Cryptosporidium* displays a strict tropism for intestinal epithelial cells (IECs), however unlike several model intestinal pathogens (such as *Salmonella*, *Listeria*, and *Toxoplasma*), *Cryptosporidium* does not breach the epithelial barrier and completes its entire lifecycle within the small intestine of a single host (22–24). It is well-established that mice deficient in IL-12p40 or IFN-γ are highly susceptible to *Cryptosporidium*, in part because these cytokines can drive innate lymphoid cell (ILC) production of IFN-γ to promote early resistance to the parasite (25–32). However, long-term control of *Cryptosporidium* is dependent on T cells, and mice and humans with primary and acquired defects in T cell function can fail to clear the parasite (25,33,34). Thus, multiple studies have highlighted that CD4^+^ Th1 responses are important for resistance to *Cryptosporidium*. Recent work has shown that type 1 conventional dendritic cells (cDC1s) are a source of IL-12 that promotes Th1 CD4^+^ responses to mediate resistance to *Cryptosporidium* (35–37). However, the ability to define the processes that promote CD4^+^ T cell responses relevant to *Cryptosporidium* has been hampered by a paucity of reagents to distinguish T cell responses to the pathogen from those activated T cells that exist in the gut at homeostasis (38–41).

In order to understand the development of parasite-specific CD4^+^ T cell responses and how these promote control of *Cryptosporidium,* transgenic parasites were engineered to express different peptide MHCII-restricted model antigens. The use of these parasite strains revealed that while *Cryptosporidium*-specific CD4^+^ T cells are primed in the mesenteric lymph node (mesLN), they only acquire full Th1 features such as T-bet and IFN-γ expression after trafficking to the gut. While cDC1s were a major source of IL-12 that is important for the generation of Th1 responses and were required for expansion and gut-homing of CD4^+^ T cells, IL-12p40 only drove local IFN-γ production and retention of CD4^+^ T cells in the gut but not early processes. These studies indicate that cDC1s provide IL-12p40-independent functions in the mesLN and -dependent functions in the gut to promote local protective CD4+ T cell responses against an enteric pathogen. Together, these studies reveal distinct roles for cDC1s in shaping CD4^+^ T cell responses to enteric infection: first to drive early expansion in the mesLN and second to drive effector responses in the gut.

## Results

### Engineering *Cryptosporidium* to express MHCII-restricted model antigens

We have recently engineered transgenic *Cryptosporidium* to express the MHCI-restricted epitope SIINFEKL at the C-terminus of the parasite effector MEDLE2 (which is secreted into the host cell cytoplasm during the intracellular stages of infection (42)) to track parasite-specific CD8^+^ T cell responses (43 Preprint). A similar approach was adopted to analyze CD4^+^ T cell responses by engineering transgenic *C. parvum* (*Cp*) to express a neomycin resistance marker and nanoluciferase (nluc) to measure fecal oocyst shedding, as well as the fluorescent protein mNeon in the parasite cytoplasm. These parasites were also modified to express either the 2W1S or the LCMV-gp61-80 (gp61) MHCII-restricted epitopes in combination with hemagglutinin (HA) as small peptide tags attached to the C-terminus of MEDLE2 (**Fig 1a, SFig 1a**). B6 mice possess a high precursor frequency of 2W1S-specific CD4^+^ T cells (44), while the gp61 peptide is the target of TCR-transgenic CD4^+^ T cells from SMARTA mice (45). To assess localization of the transgenic *Cryptosporidium* proteins, HCT8 cells were infected with either *Cp*-2W1S or *Cp*-gp61 and immunofluorescence microscopy was performed to detect the HA epitope. As shown in Fig 1b, for both constructs, the expression of mNeon (arrowed, green) provided the ability to distinguish infected from uninfected cells, while staining for HA (red) revealed that secretion of the transgenic proteins was restricted to the cytosol of infected cells (**Fig 1b**).

**Figure 1.**
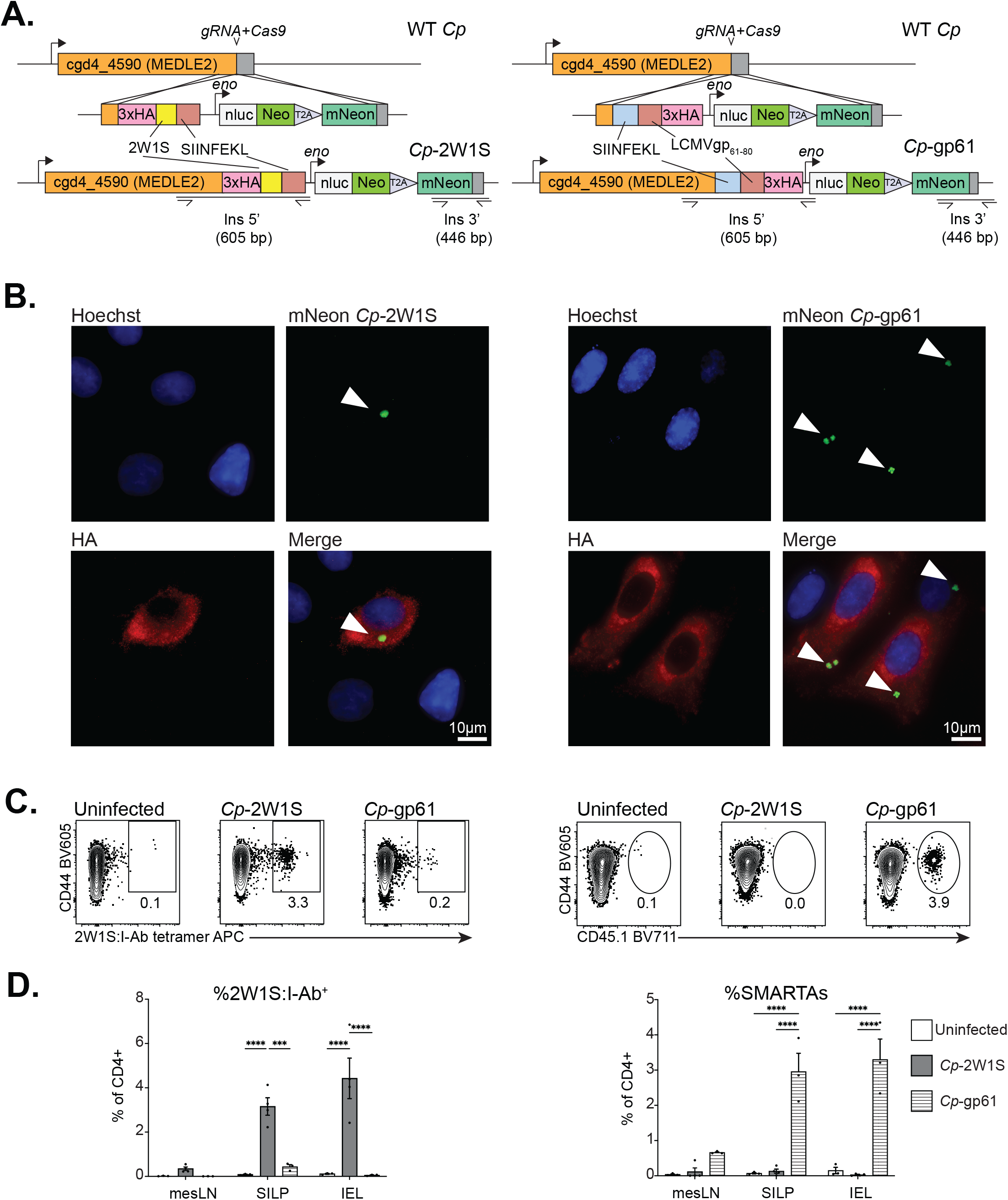
Engineering *Cryptosporidium* to express MHCII-restricted model antigens allows for identification of parasite-specific CD4^+^ T cells. **A.** Genetic construct of transgenic *Cp*-2W1S (right) or *Cp*-gp61 (left) engineered to tag the C-terminus of MEDLE2 with 3xHA-2W1S-SIINFEKL or SIINFEKL-gp61-3xHA tags. The construct also included the neomycin (Neo) selection marker and the nanoluciferase (nluc) reporter to monitor parasite burden, as well as cytoplasmic mNeon. **B.** HCT-8 cells were infected for 24 hours with 300,000 oocysts of mNeon green *Cp*-2W1S (right) or *Cp-*gp61 (left) and then stained for nuclear dye (Hoechst, blue) and HA (red). A white arrow points to the parasite within the cell. Scale bar: 10 μm. C. *Ifng*^-/-^ mice received 2×10^4^ CD45.1^+^ SMARTA T cells and were left uninfected or infected with either 1×10^4^ *Cp*-2W1S or *Cp*-gp61 oocysts and ileal draining mesenteric lymph node (mesLN), small intestinal lamina propria (SILP), or intraepithelial lymphocytes (IEL) were harvested at 10 dpi for flow cytometry. Representative flow plots show 2W1S:I-Ab+ or CD45.1^+^ SMARTA T cells in the SILP. Gated on Singlets, Live, CD19^-^, NK1.1^-^, CD90.2^+^, CD8a^-^ CD4^+^, CD44-hi, 2W1S:I-Ab^+^ or CD45.1^+^. **C.** Summary bar graphs from mesLN, SILP, and IEL of mice in (B) showing means of n=3 mice/group from 2 experiments. SMARTA T cells were CD45.1^+^CD44-hi and 2W1S-tetramer^+^ were 2W1S:I-Ab^+^CD44-hi. Statistical significance was determined in (C) by two-way ANOVA and multiple comparisons. p≤0.05, ** p≤0.01, *** p≤0.001, **** p≤0.0001.

To determine if these different parasite strains could stimulate endogenous 2W1S-tetramer^+^ CD4^+^ T cell responses or adoptively-transferred CD45.1^+^ SMARTA T cells, initial studies were performed in *Ifng*^-/-^ mice to permit robust *Cp* replication. In uninfected *Ifng*^-/-^ mice, 2W1S-tetramer^+^ and SMARTA T cells (20,000 transferred) were not detected in the draining mesenteric lymph node (mesLN), the small intestine lamina propria (SILP), or the intraepithelial lymphocyte compartment (IEL) (**Fig 1c-d**). In contrast, at 10 days post infection (dpi) *Cp-*2W1S was associated with the emergence of 2W1S-tetramer^+^ T cells but did not result in bystander activation of the SMARTA T cells. In contrast, infection with *Cp-*gp61 led to expansion of CD45.1^+^ SMARTA T cells but not the 2W1S-tetramer^+^ T cells (**Fig 1c-d**). These datasets establish that these parasite-derived MHCII-restricted model antigens lead to the antigen-specific activation of CD4^+^ T cells.

### *Cryptosporidium-*specific CD4^+^ T cells produce IFN-γ locally in the gut to control infection

Since T cell-mediated resistance to *Cryptosporidium* is associated with the production of IFN-γ, studies were performed to determine whether the response to these model antigens reflected the natural response to this pathogen. Therefore, mice in which the gene for the surface-expressed protein CD90.1 is under the control of the *Ifng* promotor (46) were infected with *Cp-*2W1S. In WT mice infected with *Cp*-2W1S, it was difficult to reliably detect 2W1S-tetramer^+^ CD4^+^ T cells, reflecting the inability of *Cp* to robustly infect immune competent WT mice (Fig 2a-b) (31). Blockade of IFN-γ during infection allowed for increased parasite burden and more reliable detection of 2W1S-tetramer^+^ cells at 10 dpi (**Fig 2a-b**) (32). In the absence of infection or without anti-IFN-γ treatment, few CD4^+^ T cells in the mesLN, SILP, or IEL expressed IFN-γ (CD90.1^+^) (**Fig 2c-d, SFig 2a**). In infected mice, when IFN-γ was blocked few (<5%) cells in the mesLN were CD90.1^+^ (**Fig 2d, SFig 2a**). In the SILP, there was minimal production of IFN-γ by the polyclonal populations, but ∼25-50% of tetramer^+^ cells were CD90.1^+^, (**Fig 2c-d**). In contrast, in the IEL, ∼50-75% of tetramer^+^ cells were CD90.1^+^, with similar percentages in polyclonal CD4^+^ IELs (**Fig 2c-d**).

**Figure 2.**
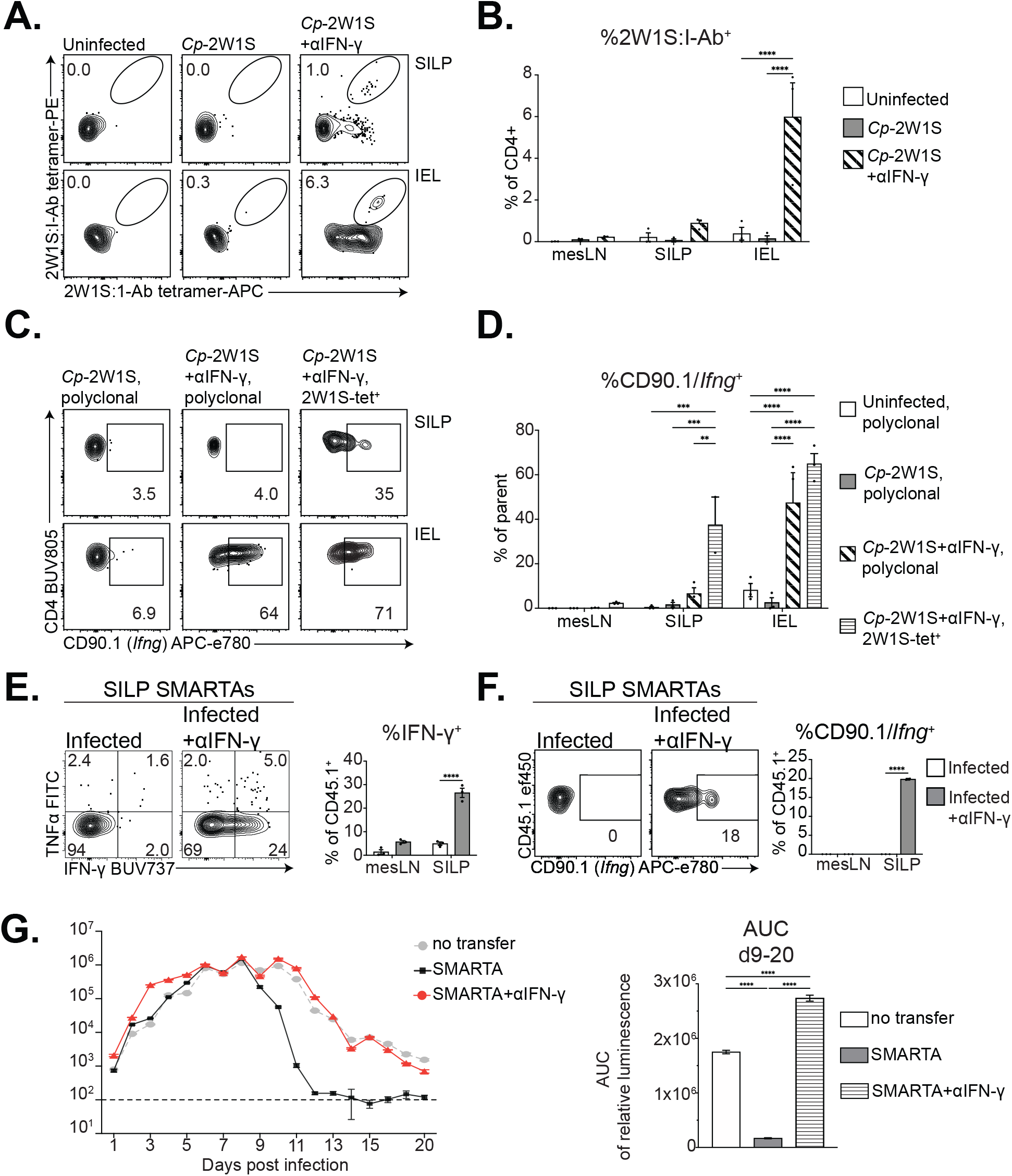
*Cryptosporidium*-specific CD4^+^ T cells produce IFN-γ locally to protect against infection. **A-D.** IFN-γ-CD90.1 reporter mice were left untreated or treated with 1mg/mouse of αIFN-γ 1 day prior to infection and 2, 5, and 8 dpi with 10^4^ *Cp*-2W1S. mesLN, SILP, and IEL were harvested at 10 dpi for flow cytometry. Data shown are from 1 experiment representative of 2 independent experiments. **A.** Representative flow plots from the SILP and IEL of CD4^+^ T cells stained with 2W1S:I-Ab tetramer in PE and APC showing detection of tetramer+ cells when mice are infected and treated with IFN-γ. Gating: Singlets, Live^+^, CD19^-^, NK1.1^-^, EpCAM^-^, CD90.2^+^, CD8a^-^, CD4^+^. **B.** Quantification/summary of (A) and tetramer (PE and APC double positive) staining from mesLN, SILP, and IEL. **C.** Representative flow plots showing CD90.1 expression in polyclonal CD4^+^ T cells (first two columns) or 2W1S:I-Ab tetramer^+^ cells (last column) in SILP and IEL of infected mice untreated (first column) or treated with αIFN-γ (last two columns). Gating: Singlets, Live^+^, CD19^-^, NK1.1^-^, EpCAM^-^, CD90.2^+^, CD8a^-^, CD4^+^, 2W1S:I-Ab^+^. **D.** Quantification/summary of (C) showing the percent of cells that were CD90.1^+^ from the following groups: polyclonal CD4^+^ T cells from uninfected mice (white), polyclonal CD4^+^ T cells from untreated mice infected with *Cp*-2W1S (gray), polyclonal CD4^+^ T cells from αIFN-γ-treated mice infected with *Cp*-2W1S (white with diagonal stripes), or 2W1S:I-Ab tetramer^+^ CD4^+^ T cells from αIFN-γ-treated mice infected with *Cp*-2W1S (white with horizontal stripes). **E-F.** WT B6 mice received 2×10^4^ CD90.1/*Ifng* reporter SMARTA T cells 1 day prior to infection with 5×10^4^ ma*Cp*-ova-gp61 and were left untreated or treated with 1mg/mouse of αIFN-γ 1 day prior to infection and 2, 5, and 8 dpi. At 10 dpi mesLN and SILP were harvested for flow cytometry and cells were stimulated with exogenous gp61 peptide for 3 hours followed by intracellular cytokine staining and flow cytometry. Data shown are from 1 experiment representative of 2 independent experiments. **E.** Left: representative flow plots and from the SILP showing IFN-γ and TNF-α expression in SMARTA T cells after peptide stimulation. Gating: Singlets, Live^+^, CD19^-^, NK1.1^-^, EpCAM^-^, CD90.2^+^, CD8a^-^, CD4^+^, CD45.1^+^. Right: summary of showing the percentage of SMARTA cells from the SILP or mesLN staining IFN-γ+ after peptide stimulation. **F.** Left: representative flow plots from the SILP showing expression of CD90.1 as a reporter of *Ifng* expression on SMARTA T cells after peptide stimulation. Gating: Singlets, Live^+^, CD19^-^, NK1.1^-^, EpCAM^-^, CD90.2^+^, CD8a^-^, CD4^+^, CD45.1^+^. Right: summary showing the percentage of SMARTA T cells from the SILP or mesLN staining for CD90.1^+^ after peptide stimulation. **G.** PBS or 10^6^ SMARTA T cells were transferred into *Ifng*^-/-^ mice 1 day prior to infection with 10^4^ *Cp*-gp61 and feces was analyzed by nanoluciferase for parasite burden (relative luminescence) over time. For some mice receiving SMARTA T cells, mice also received 1mg/mouse of αIFN-γ blocking antibody 1 day prior to infection and 2, 5, and 8 dpi. Area under the curve analysis was performed for each treatment for 9-20 dpi. Graphs shown are from 1 experiment, representative of 2 independent experiments. Statistical significance was determined by two-way ANOVA and multiple comparisons. p≤0.05, ** p≤0.01, *** p≤0.001, **** p≤0.0001.

Likewise, SMARTA T cells were isolated from the mesLN and SILP of WT mice infected with gp61-expressing parasites, stimulated with gp61 peptide *ex vivo*, and subjected to intracellular staining for IFN-γ. A cohort of mice was treated with anti-IFN-γ during infection to increase parasite burden. Regardless of anti-IFN-γ treatment, few SMARTA T cells in the mesLN produced IFN-γ after peptide restimulation (**Fig 2e**, **SFig 2b**). Among SILP SMARTA T cells from untreated mice, <5% of cells produced IFN-γ, whereas 20-30% of SMARTAs from anti-IFN-γ-treated IFN-γ^+^ (**Fig 2e**). IFN-γ production could also be detected in IFN-γ reporter SILP but not mesLN SMARTA T cells that expressed CD90.1 under the control of the *Ifng* promotor (**Fig 2f**). These data collectively indicate that the ability of *Cryptosporidium*-specific CD4^+^ T cells to produce IFN-γ (even during restimulation with exogenous peptide) is restricted to the SILP and IEL and does not occur in the mesLN.

In order to determine whether SMARTA CD4^+^ T cell-derived IFN-γ is protective, *Ifng*^-/-^ mice were infected with *Cp-*gp61. One day prior to infection, a cohort received 10^6^ IFN-γ-sufficient at CD4*^+^* T cells. In *Ifng*^-/-^ mice that did not receive T cells, infection peaked at ∼d9 and oocyst shedding subsequently decreased though mice remained chronically infected (**Fig 2g**). In mice that received IFN-γ-sufficient SMARTA T cells, infection also peaked at ∼d9, but oocyst shedding dropped faster and fell below the limit of detection (**Fig 2g**). This protective effect of the SMARTA T cells was abolished in mice treated with anti-IFN-γ (**Fig 2g**). These datasets highlight the utility of this transgenic system and reveal a protective effect for CD4^+^ T cell-derived IFN-γ whose production is limited to the gut.

### *Cryptosporidium*-specific CD4^+^ T cells express distinct activation states associated with Th1 responses and mucosal tissue residency

Next, WT mice were utilized to study CD4^+^ T cell responses in an immune-competent setting. Because conventional *Cp* does not readily infect WT mice, a mouse-adapted strain of mCherry-expressing *C. parvum* (ma*Cp*) (32) was engineered to express model antigens. As the ma*Cp* strain was previously engineered with neomycin resistance to drive mCherry and nluc expression, a second marker was introduced to confer resistance to the recently-described bicyclic azetidine BRD7929 that targets *Cp* phenylalanine tRNA synthetase (pheRS) (47,48). Hence, the *Cp* pheRS locus was modified to alter leucine at position 482 to valine, conferring resistance to BRD7929 as previously described (49). Model-antigen-tagged MEDLE2 was introduced under the *Cp*-*enolase* promotor immediately following the transgenic pheRS^R^, such that BRD7929-resistant parasites (ma*Cp*-ova-gp61) would express ectopic MEDLE2-SIINFEKL-gp61 (**SFig 3a-b**). Infection of WT mice with ma*Cp*-ova-gp61 combined with adoptive transfer of SMARTA T cells and high-parameter flow cytometry provided the opportunity to compare polyclonal and parasite-specific CD4^+^ T cell responses against *Cryptosporidium* in WT mice. This comparison allowed for assessment of CD4^+^ T cell expression of transcription factors (TFs) for Th lineages (T-bet for Th1, RORγΤ for Th17, and Foxp3 for Treg) and surface markers associated with Th1 cells (CXCR3, SLAM, IL-18Ra), antigen-experience/activation (CD44, CD69, CD40L, Ly6A/E), and mucosal association (LPAM-1, CD103). UMAP projection of aggregated CD4^+^ T cells from all tissues (SMARTAs and non-SMARTAs from the mesLN, SILP, and IEL) and conditions (uninfected and infected) was performed. A comparison of CD4^+^ T cells by tissue highlighted that mesLN and IEL CD4^+^ T cells occupied distinct regions in UMAP space while those in the SILP resembled both the mesLN and IEL (**Fig 3a**). When X-Shift unbiased clustering analysis of the aggregated CD4^+^ T cells was performed, 11 clusters were apparent (**Fig 3b-c, SFig 3c**). Cluster 1 consisted mainly of CD4^+^ T cells from the mesLN that were LPAM-1-hi (**Fig 3b, SFig 3e**). Th1 CD4^+^ T cells could be identified by high T-bet expression in clusters 2-7, while Th17 associated with RORγT expression was limited to cluster 8, and Foxp3^+^ Tregs could be found in clusters 9-11 (**Fig 3b, SFig 3e**). Further analysis of the T-bet^+^ clusters showed that the majority of clusters 3-6 was composed of CD4^+^ T cells from the SILP and IEL of infected mice that increased in frequency during *Cryptosporidium* infection (**Fig 3d-e, SFig 3d**).

**Figure 3.**
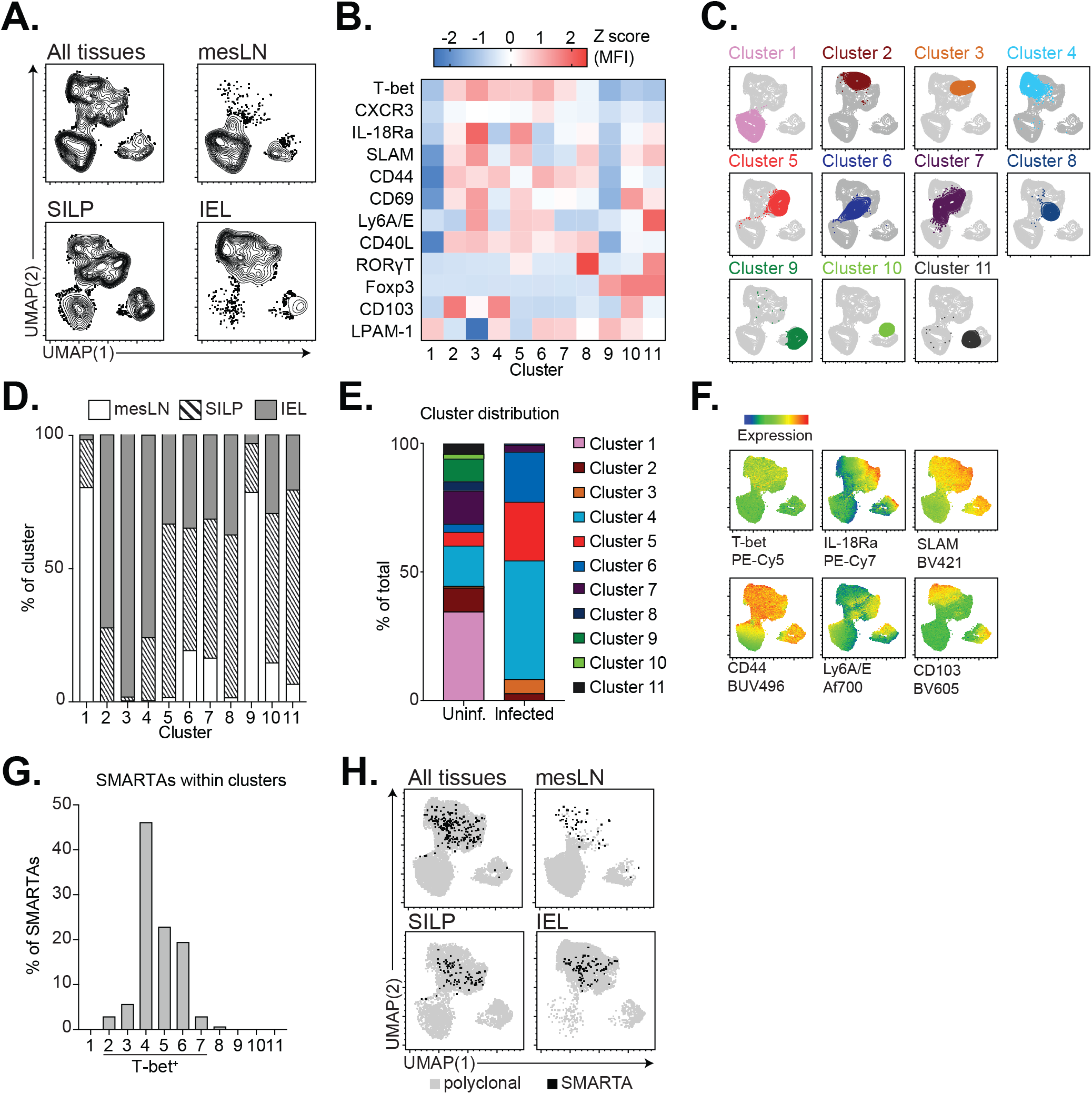
*Cryptosporidium*-specific CD4^+^ T cells resemble polyclonal T-bet^+^ cells in the gut. **A.** WT B6 mice received 2×10^4^ SMARTA T cells 1 day prior to infection with 5×10^4^ ma*Cp*-ova-gp61 oocysts and mesLN, SILP, and IEL were harvested at 10 dpi for flow cytometry. **A.** UMAP of mesLN, SILP, and IEL CD4^+^ T cells (Singlets, Live^+^, CD8a-EpCAM-NK1.1^-^CD19^-^CD4^+^TCRβ^+^) based on surface expression of the following markers: IL-18Ra, SLAM, CXCR3, T-bet, RORγT, Foxp3, CD40L, CD44, CD69, CD103, Ly6A/E, and LPAM-1. **B.** X-shift cluster analysis of concatenated CD4^+^ T cells from all tissues revealed 11 clusters. Heatmap of the Z-scores of surface marker expression by X-shift cluster. **C.** Overlays of each cluster onto the UMAP from Figure 3A**. D.** The percentage of cells in each cluster coming from mesLN, SILP or IEL. **E.** The percentage of total CD4^+^ T cells represented by each cluster in uninfected or infected mice. **F.** UMAP from Figure 3A colored by expression levels of each marker. **G.** Percentage of SMARTA T cells that fell into each cluster, with the total across all clusters equalling 100%. **H.** UMAP showing where SMARTA T cells fell in the UMAP (black dots).

Because SMARTA T cells were only detected in these tissues during infection (**Fig 1b-c**), this provided an opportunity to compare the *Cryptosporidium*-induced SMARTA CD4^+^ T cells to polyclonal responses. In UMAP space, SMARTA T cells fell into a CD44-hi region near highly activated Foxp3^+^ cells in all tissues (**Fig 3f-h**). However, the majority of SMARTA T cells were present in T-bet^+^ clusters and were most represented in clusters 4 and 5 (**Fig 3g**). Cluster 4 was CD103^+^ while cluster 5 was associated with high levels of IL-18Ra, SLAM, and the activation marker Ly6A/E. Across all T-bet^+^ clusters, CD103 and Ly6A/E were inversely correlated and cells occupied different regions in UMAP space (**Fig 3b, f**).

This analysis indicates that the infection-induced activation of the SMARTA T cells mirrors the processes associated with the endogenous polyclonal responses. Therefore, this system described above was used to understand how SMARTA T cell expression of lineage-defining transcription factors (TFs; T-bet, GATA3, RORγΤ, Foxp3) and Tfh markers (Bcl-6, PD-1, CXCR5) varied across tissues. In the mesLN of uninfected mice the majority (>80%) of (non-SMARTA) CD4^+^ T cells expressed no lineage-defining TFs (T-bet^-^GATA3^-^RORγΤ^-^Foxp3^-^; henceforth referred to as TF^-^), and therefore appeared uncommitted to any one Th subset. At 10 dpi, despite an increased number of CD4^+^ T cells in the mesLN, the majority (∼80%) of polyclonal CD4^+^ T cells in this tissue remained TF^-^ (**Fig 4a-b, SFig 4a-b**). However, all SMARTA T cells in the mesLN of infected mice were antigen experienced (∼100% CD44^+^) and ∼30-40% were T-bet^+^ (**Fig 4a-b**).

**Figure 4.**
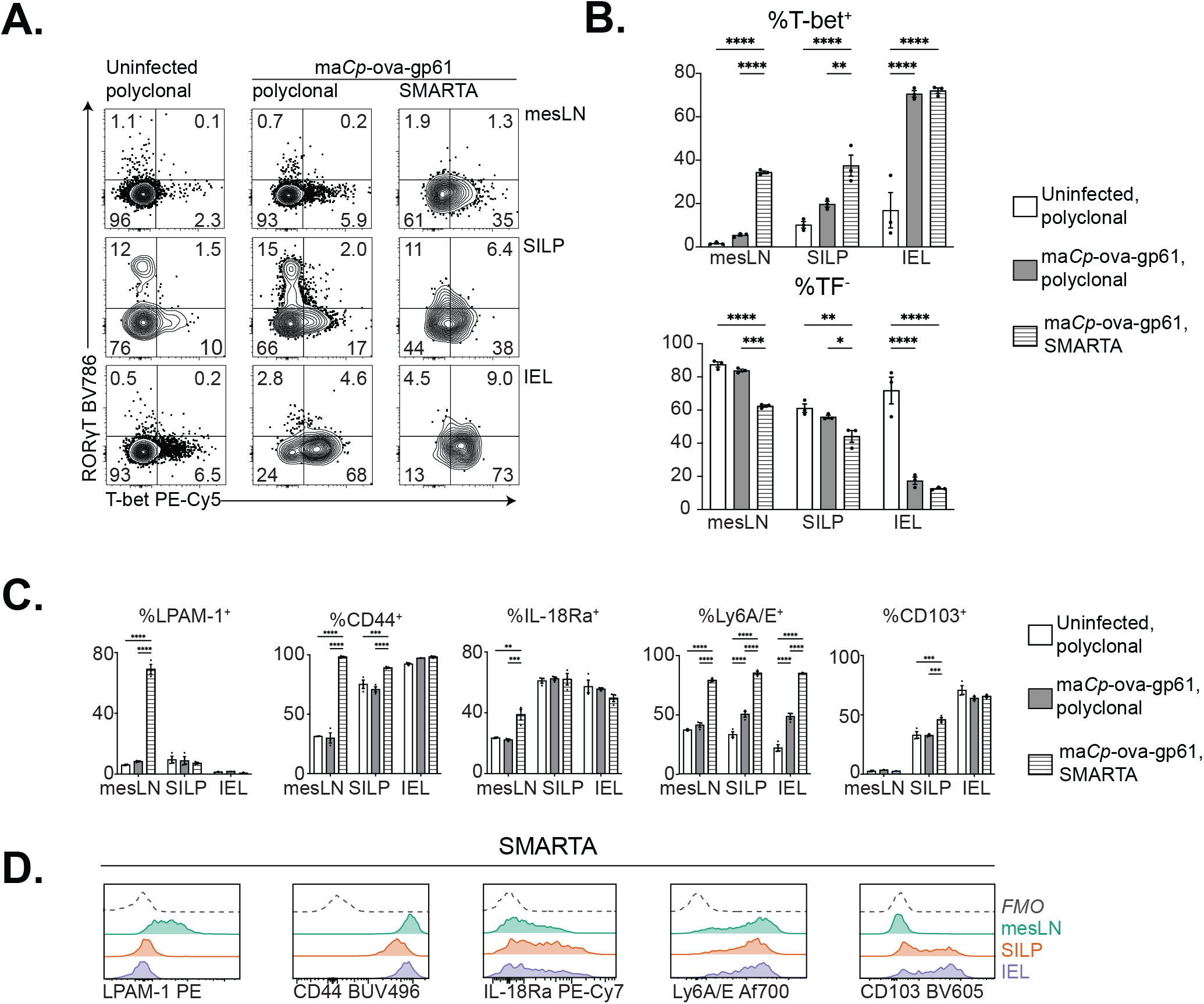
Progressive Th1 skewing of *Cryptosporidium*-specific CD4^+^ T cells. WT B6 mice received 2×10^4^ SMARTA T cells 1 day prior to infection with 5×10^4^ ma*Cp*-ova-gp61 oocysts and mesLN, SILP, and IEL were harvested at 10 dpi for flow cytometry. **A.** Representative flow plots of T-bet vs. RORγΤ in polyclonal CD4^+^ T cells from uninfected mice, polyclonal CD4^+^ T cells from infected mice, or SMARTA T cells (CD45.1^+^) from infected mice organized by tissue (rows). Gating: Singlets, Live^+^, CD19^-^, NK1.1^-^, EpCAM^-^, CD90.2^+^, CD8a^-^, CD4^+^, CD45.1^+^. **B.** Percentage of T-bet^+^, or percentage negative for all lineage-defining TFs (T-bet, RORγΤ, Foxp3, GATA3; TF^-^) in polyclonal CD4^+^ T cells from uninfected mice (white), polyclonal CD4^+^ T cells from infected mice (gray), or SMARTA T cells (horizontal lines). **C.** Percentage of cells positive for LPAM-1, CD44, IL-18Ra, Ly6A/E, or CD103 of polyclonal CD4^+^ T cells from uninfected mice, infected mice, or SMARTA T cells colored as in (B). **D.** Histograms of protein expression by flow cytometry among SMARTA T cells from the mesLN (teal), SILP (orange) or IEL (purple)). Gating: Singlets, Live+, CD8a^-^EpCAM^-^ NK1.1^-^CD19^-^CD4^+^TCRβ^+^CD45.1^+^. For A-B, representative of 2 independent experiments; for C-D, one independent experiment. n=3 mice/group. Statistical significance was determined by two-way ANOVA and multiple comparisons. p≤0.05, ** p≤0.01, *** p≤0.001, **** p≤0.0001.

In the SILP of uninfected mice, distinct Treg (Foxp3^+^), Th17 (RORγT^+^), and Th1 (T-bet^+^) populations could be identified, although 60% of CD4^+^ T cells here were TF^-^ and no Th2 cells (GATA3^+^) could be identified (**Fig 4a-b, SFig 4a-b**). While infection did not lead to a significant change in the percentage of SILP polyclonal CD4^+^ T cells that were TF^-^, ∼40% of SMARTA T cells were T-bet^+^ compared to ∼20% of the polyclonal CD4^+^ T cells (**Fig 4a-b**). In the IEL of uninfected mice, 60% of CD4^+^ T cells were TF^-^ (similar to the mesLN and SILP) but, after infection the percentage of CD4^+^ T cells that were TF^-^ in the IEL decreased to <20% (**Fig 4b**). This was accompanied by a significant increase in the number and relative percentage of T-bet^+^ Th1 CD4^+^ T cells in the IEL (**Fig 4a-b, SFig 4a**). Similarly, >75% of SMARTA T cells in the IEL expressed T-bet (**Fig 4a-b**) and also expressed IL-18Ra, SLAM, CXCR3, Ly6A/E and CD69 (**Fig 4c-d, SFig 4d-e**). The only marker significantly higher in mesLN SMARTAs compared to in the gut was LPAM-1 (**Fig 4c-d**), consistent with its role in trafficking to the gut and subsequent downregulation. Additionally, a small percentage (3-4%) of SMARTA T cells in the mesLN were CXCR5-hi, PD-1-hi, and Bcl6-hi— suggesting some antigen-specific cells were Tfh (**SFig 4f-g**). This ability to focus on the SMARTA T cell response highlights that the parasite-specific CD4^+^ T cell response to *Cryptosporidium* is characterized by progressive changes in activation status as T cells transit from priming in the mesLN to effector functions in the IEL.

### *Cryptosporidium* infection induces differential activation of cDC1s required for expansion and gut homing of CD4^+^ T cells

Although DCs and IL-12 are important for resistance to *Cryptosporidium* (35,36), the role of different DC subsets on CD4^+^ T cell responses to *Cryptosporidium* is unclear. Therefore, DCs in the mesLN and SILP were profiled for activation markers and IL-12p40 production in order to understand how infection alters these accessory cells. At 4 dpi, the numbers of cDC1s and cDC2s in the mesLN and SILP were not significantly changed (**SFig 5a**). Additionally, cDC2s showed little change in expression of activation markers (CD40, CD86, MHCII) or IL-12p40 expression at 4 dpi (**Fig 5a-b, SFig a-c**). In contrast, cDC1s showed increased expression of CD40 in the mesLN and SILP, and up-regulation of CD86 and MHCII in the SILP (**Fig 5a-b**). Regardless of infection status, cDC1s in the mesLN produced low levels of IL-12p40 (**Fig 5c, SFig 5a**) (50). However, after infection, cDC1s showed a marked upregulation of IL-12p40 in the SILP (**Fig 5c-d**). Thus, locally-activated cDC1s within the SILP are a major source of IL-12p40 during *Cryptosporidium* infection.

**Figure 5.**
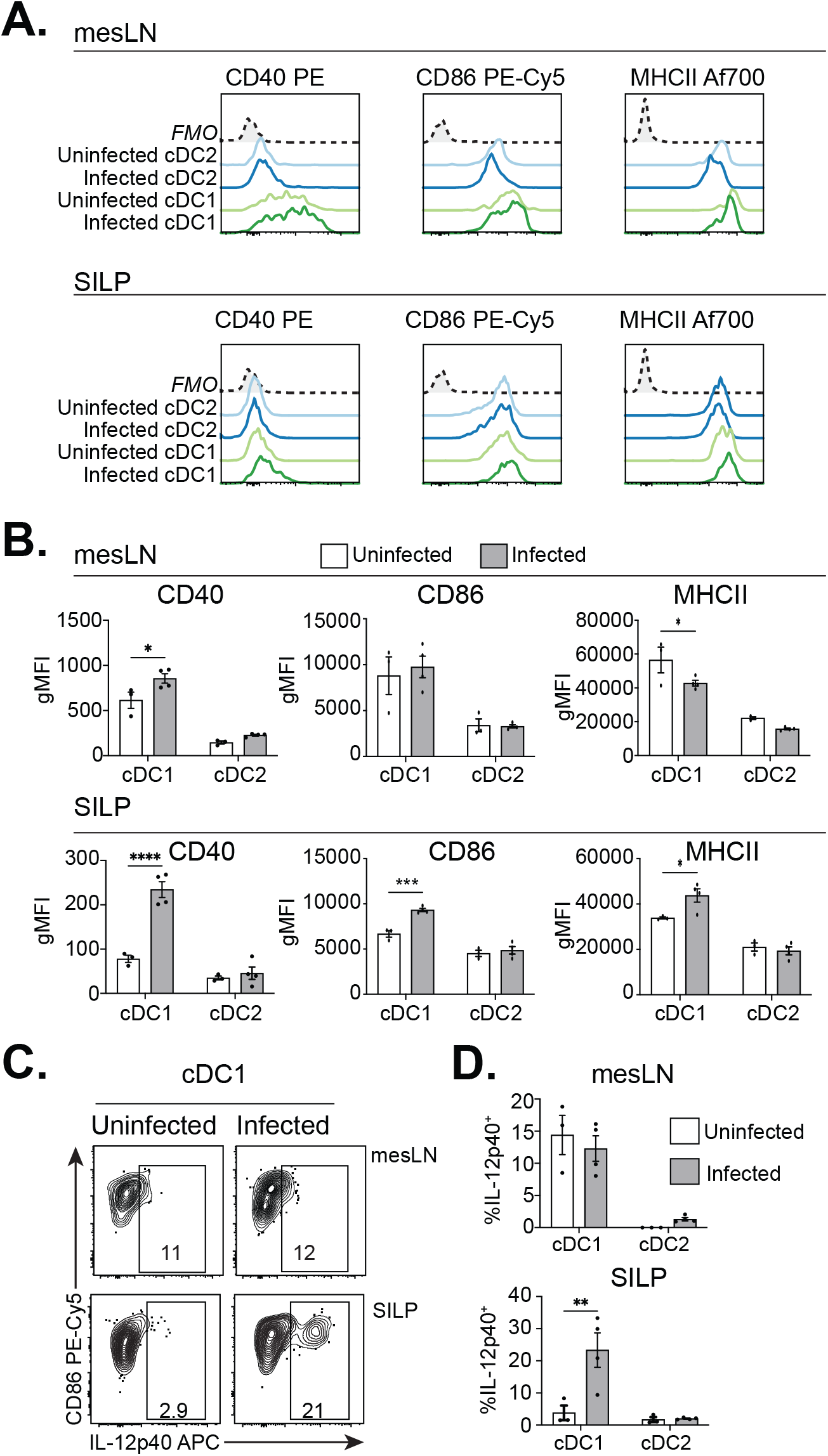
IL-12p40 is produced in the gut by activated cDC1s. **A-D.** WT B6 mice were left uninfected or infected with 5×10^5^ ma*Cp* oocysts. At 4 dpi mesLN and SILP cells were isolated, plated with brefeldin A (BFA) for 6 hours, and analyzed by flow cytometry for surface markers and IL-12p40 expression. Plotted are histograms from mesLN or SILP (A), summary in (B); IL-12p40 flow plots from the mesLN and SILP in (C) summarized in (D) showing selective induction of IL-12p40 in cDC1s in the SILP of infected mice. Data shown are from 1 experiment representative of 2 independent experiments, 3-4 mice per group. Gating: Singlets, Live+, CD3e^-^, NK1.1^-^, EpCAM^-^, B220^-^, CD19^-^, CD90.2^-^, CD64^-^, MHCII-hi, CD11c^+^, CD26^+^, Ly6C^-^, XCR1^+^ (cDC1s) or SIRPα^+^ (cDC2s). Gating is based on FMOs taken from SILP for all colors except IL-12p40, for which positivity is based on samples from each tissue plated without BFA. Statistical significance was determined by two-way ANOVA and multiple comparisons. p≤0.05, ** p≤0.01, *** p≤0.001, **** p≤0.0001.

To examine the contribution of cDC1s to CD4^+^ T cell responses to *Cryptosporidium*, 1×10^6^ SMARTA T cells that express Nur77-GFP as a reporter of TCR activation (51) were labeled with CellTrace Violet (CTV). One day prior to infection, these cells were transferred into WT control mice and mice that lack cDC1s (*Irf8+32*^-/-^). At 1 dpi with ma*Cp*-ova-gp61, in WT mice few SMARTA T cells could be found in the mesLN, and by 4 dpi small numbers of Nur77^+^ SMARTAs were present but had not yet proliferated (**Fig 6a**). By 6 dpi, >90% of SMARTA T cells had divided, and those cells that had undergone >5 rounds of division had up-regulated CXCR3 and the gut homing receptor LPAM-1 (**Fig 6b-e**). Similar to WT mice, in *Irf8+32*^-/-^ mice few SMARTA T cells could be found in the mesLN on 1 dpi, where small numbers of Nur77^+^ SMARTAs were present but had not yet proliferated (**Fig 6a**). By 6 dpi in the mesLN of *Irf8+32*^-/-^ mice, a lower percentage of SMARTAs had divided than in WT mice (80% in *Irf8+32*^-/-^ mice compared to >90% in WT, **Fig 6a-b**). Strikingly in contrast to WT mice, in *Irf8+32*^-/-^ mice, the most-divided SMARTA T cells had reduced ability to express LPAM-1 as well as CXCR3 (**Fig 6c-e**). Consistent with this observation, despite elevated parasite burden (data not shown), at 10 dpi *Irf8+32*^-/-^ mice had reduced numbers of SMARTA T cells in the mesLN and an absence of these cells in the SILP and IEL (**Fig 6f-g**). Additionally at this timepoint, while in WT mice SMARTA T cells expressed IL-18Ra in the mesLN, in *Irf8+32*^-/-^ mice cells did not express this receptor (**Fig 6h-i**). Together, these data suggest CD4^+^ T cells responding to *Cryptosporidium* do not require cDC1s for priming but instead rely on these cells for early signals leading to optimal proliferation, cytokine/chemokine receptor expression, and homing to the gut.

**Figure 6.**
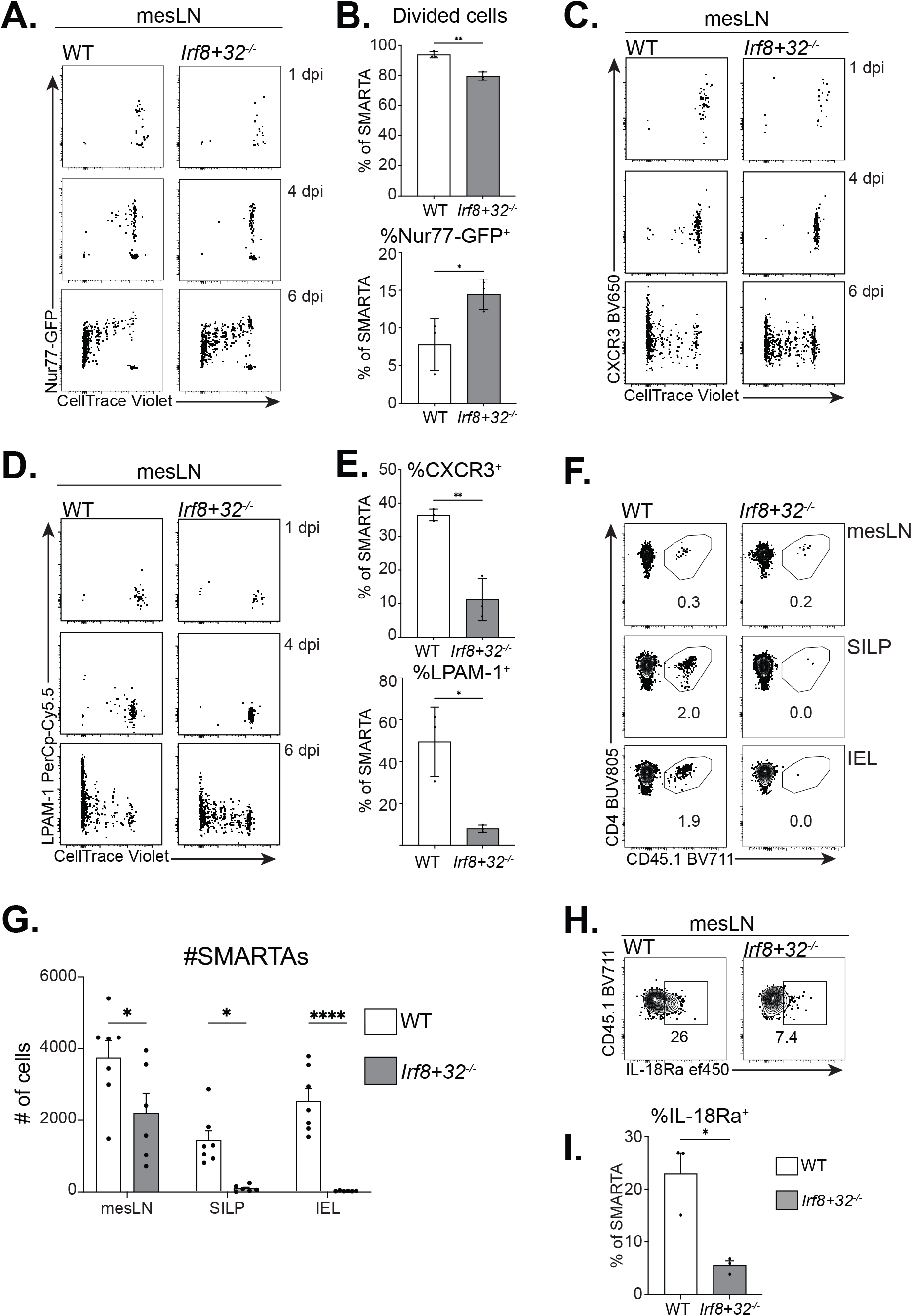
cDC1s are required for expansion and gut homing of *Cryptosporidium*-specific CD4^+^ T cells. **A-E.** 1 day prior to infection with 5×10^4^ ma*Cp*-ova-gp61 oocysts, 1×10^6^ CD45.1^+^ Nur77-GFP reporter SMARTA T cells were labeled with CellTrace Violet (CTV) and transferred into WT B6 mice or age/ sex-matched *Irf8+32*^-/-^ mice. Mice were sacrificed at 1, 4, and 6 dpi and priming of SMARTA T cells was interrogated using flow cytometry. **A-B.** Representative flow plots from the mesLN of infected mice at 1, 4, and 6 dpi pre-gated on SMARTA T cells showing Nur77-GFP expression compared to cell division, with summary in (B). **C-E** Representative flow plots from mesLN SMARTAs showing CellTrace Violet versus CXCR3 (C) and LPAM-1 (D) with summary in (E). For A-E, data is from 1 experiment representative of 2 independent experiments. Gating for SMARTAs in A-E: Singlets, Live^+^, NK1.1^-^, CD19^-^, EpCAM^-^, CD90.2^+^, CD8a^-^, CD4^+^, TCR Vβ8.3^+^, CD45.1^+^. **F-I.** 1 day prior to infection with 5×10^4^ ma*Cp*-ova-gp61 oocysts, 2×10^4^ CD45.1^+^ SMARTA T cells were transferred into WT B6 mice or age/sex-matched *Irf8+32*^-/-^ mice. At 10 dpi, mesLN, SILP, and IEL were isolated and analyzed by flow cytometry. Data are representative of 2 independent experiments, n=3 mice/group. **F.** Representative flow plots showing presence or absence of SMARTA T cells in the mesLN (top), SILP (middle), or IEL (bottom). Plots are from 1 experiment representative of 3 independent experiments. **F.** Summary of (G). **H-I.** Percentage of mesLN SMARTA T cells staining positive for IL-18Ra with representative flow plots in (H) and summary in (I). Gating for SMARTAs in G-I: Singlets, Live^+^, NK1.1^-^, CD19^-^, EpCAM^-^, CD90.2^+^, CD8a^-^, CD4^+^, CD44-hi, CD45.1^+^.

### IL-12p40 is dispensable for priming but is required for CD4^+^ T cell production of IFN-γ in the gut

To determine the role of IL-12p40 on the CD4^+^ T cell response, SMARTA CD4^+^ T cells were profiled using in vivo blockade of IL-12p40. IL-12p40 blockade led to a marked increase in parasite burden (**Fig 7a**). Blockade of IL-12p40 did not impact the ability of SMARTA CD4^+^ T cells to traffic to the SILP or IEL (**Fig 7b**). However, IL-12p40 blockade did reduce the absolute number of SMARTA T in the IEL, with similar results in *Il12b^-^*^/-^ mice (**Fig 7b-c, SFig 7a-b**). Blockade of IL-12p40 also led to a reduction in expression of T-bet and IL-18Ra in SMARTA T cells (**Fig 7d-g**). Because IL-18 and IL-12 can drive CD4^+^ T cell IFN-γ production (52), the influence of these cytokines on IFN-γ expression by SMARTA T cells was assessed. SMARTA T cells expressing the surface protein CD90.1 under the control of the *Ifng* promotor were transferred into *Ifng*^-/-^ mice (in order to remove confounding effects of IL-12p40 or IL-18 on innate cell production of IFN-γ) 1 day prior to infection with *Cp*-gp61. Mice were treated with isotype control, anti-IL-18, anti-IL-12p40, or anti-IL-18 plus anti-IL-12p40 1 day prior to infection as well as 2, 5, and 8 dpi. Mice were sacrificed at 10 dpi and expression of CD90.1 and IL-18Ra on SMARTA T cells was assessed by flow cytometry. In control mice roughly ∼40% of SMARTA T cells in the SILP and ∼60% of SMARTA T cells in the IEL were CD90.1^+^, and treatment with anti-IL-18 did not alter these responses (**Fig 7h-i**). In contrast, blockade of IL-12p40 led to a marked reduction in the percentage of SMARTA T cells that were CD90.1^+^ and a reduction in IL-18Ra expression (**Fig 7h-i, SFig 7c**). Blockade of IL-18 alone or in combination with anti-IL-12p40 did not lead to further reductions in the percentage of cells that were CD90.1^+^ (**Fig 7h-i**). These data suggest that IL-12p40 is not required for initial priming, expansion and gut-homing of *Cryptosporidium* CD4^+^ T cells but is required for their accumulation in the gut and local expression of T-bet, IL-18Ra, and production of IFN-γ.

**Figure 7.**
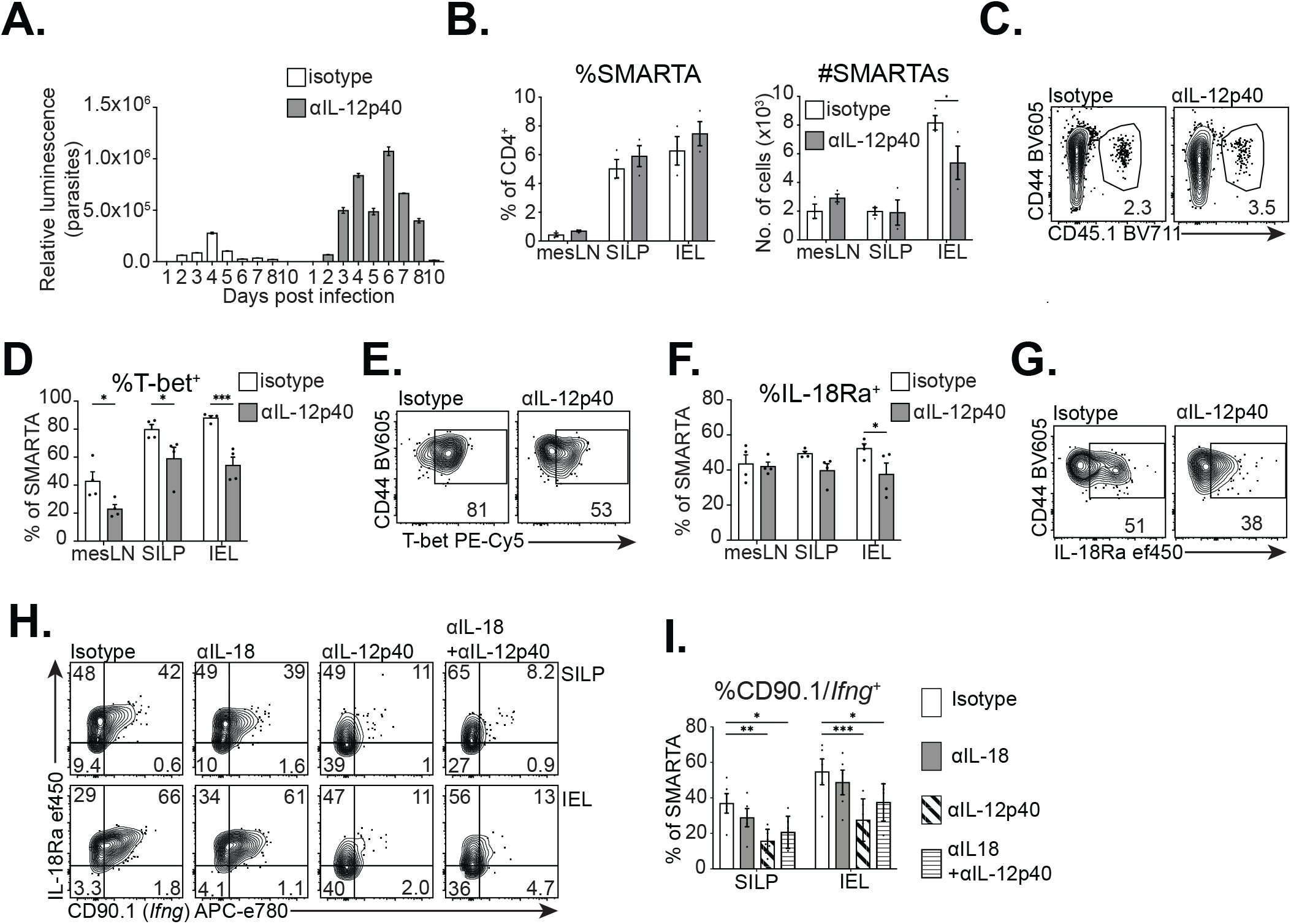
IL-12p40 is not required for gut-homing but is required for gut CD4^+^ T cell accumulation and induction of Th1 functions in *Cryptosporidium-*specific CD4^+^ T cells. **A-G.** 1 day prior to infection with 5×10^4^ ma*Cp*-ova-gp61 oocysts, 2×10^4^ CD45.1+ SMARTA T cells were transferred into WT B6 mice that were treated with isotype control (rat IgG2a) or anti-IL-12p40 (1mg/mouse on d-1, or 2, 5, 8 dpi). At 10 dpi, mesLN, SILP, and IEL were isolated and analyzed by flow cytometry. Data shown in A-G is 1 experiment representative of 2 independent experiments, n=3-4 mice/group. **A.** Infection was monitored by nanluciferase of feces. **B-C.** Percentage of SMARTA T cells among CD4^+^ T cells and absolute numbers of SMARTA T cells from each treatment with representative flow plots from the SILP in (C). **D-E.** Percentage of SMARTA T cells staining T-bet+, with representative flow plots from the SILP in (E). **F-G**. Percentage of SMARTA T cells staining for IL-18Ra, with representative flow plots from the SILP in (G). **H-I**. 1 day prior to infection with 10^4^ *Cp*-gp61, 5×10^4^ CD45.1^+^ SMARTA T cells that expressed CD90.1 as a reporter for *Ifng* expression were transferred into *Ifng*^-/-^ mice that were treated with isotype control (rat IgG2a), anti-IL-12p40, anti-IL-18, or anti-IL12p40+anti-IL-18 (1mg/mouse on d-1, or 2, 5, 8 dpi). Mice were also treated with 1mg/mouse anti-IFN-γ on 3 and 7 dpi. At 10 dpi, mesLN, SILP, and IEL were isolated and analyzed by flow cytometry and cells were stained for CD90.1 to assess *in vivo* IFN-γ production. Data shown for K is pooled from 2 independent experiments, n=3 mice/group/experiment. Gating for SMARTAs in B-K: Singlets, Live+, NK1.1^-^, CD19^-^, EpCAM^-^, CD90.2^+^, CD8a^-^, CD4^+^, CD44-hi, CD45.1^+^. Statistical significance was determined by two-way ANOVA and multiple comparisons. p≤0.05, ** p≤0.01, *** p≤0.001, **** p≤0.0001.

## Discussion

Although it has been accepted for several decades that CD4^+^ T cells are essential for clearance of *Cryptosporidium* (53,54), there are many fundamental questions about how these protective lymphocytes are generated and mediate resistance to this common enteric infection. The absence of well characterized endogenous MHCII-restricted *Cryptosporidium* antigens has made it a challenge to distinguish parasite-specific CD4^+^ T cell responses from the many T cells in the gut that possess an “activated but resting” phenotype at baseline (38–41). To overcome these obstacles, we took advantage of the recent development of parasite transgenesis (55) to engineer *Cryptosporidium* to express MHCII-restricted model antigens. When combined with MHCII-tetramers and SMARTA TCR transgenic T cells these parasites allowed for the identification of CD4^+^ T cells that have responded to *Cryptosporidum*-derived antigens. In accord with current models, these CD4^+^ T cell responses to *Cryptosporidium* were dominated by the production of IFN-γ in the gut that was sufficient to mediate parasite clearance. This tool set also allowed us to study the initial priming of these CD4^+^ T cells and unexpectedly highlighted that recently activated SMARTA T cells in the mesLN were unable to make IFN-γ and only acquired the ability to produce IFN-γ in the SILP and IEL, which was associated with expression of T-bet. While cDC1s were not required for initial priming of the SMARTA CD4^+^ T cells, these accessory cells did have a dual role in CD4^+^ T cell responses: first in the mesLN to drive early expansion and gut homing, and later, at the site of infection where cDC1 production of IL-12p40 stimulated CD4^+^ T cell IFN-γ production, T-bet expression, and tissue retention. Thus, *Cryptosporidium*-specific CD4^+^ T cells are primed in the mesLN but require signals within the gut— including IL-12p40—to achieve full effector capacity.

In other systems, early IL-12 has been shown to be important in the initial development of Th1 responses (56,57), but whether it is needed at later time points for Th1 effector responses depends on the experimental system (58–61). One possible explanation that might influence the degree of Th1 commitment in CD4^+^ T cells after priming is whether the pathogen is restricted to a barrier site like the gut, skin, or lungs, or is one that disseminates more widely. For systemic pathogens like *Toxoplasma gondii*, parasite dissemination leads to the early presence of parasites in lymphoid tissues and local production of IL-12 that results in the rapid commitment to the Th1 phenotype (58,62). When this occurs following oral infection with *T. gondii*, these heightened CD4^+^ T cell responses can result in an IFN-γ-mediated lethal ileitis (63). For *Cryptosporidium*, there is no evidence of immunopathology mediated by CD4^+^ T cells and perhaps a lack of commitment to a Th1 effector program during priming ensures that parasite-specific CD4^+^ T cells provide highly localized production of IFN-γ and thereby mitigate IFN-γ-mediated immunopathology. This is similar to the concept that CD4^+^ T cell plasticity allows them to respond to contextual signals to tailor their function to the different infection. Indicative of this plasticity, influenza-specific Th1 CD4^+^ Trm in the lung exposed to Th2-inducing signals will down-regulate T-bet and IFN-γ production (64), while the ability of conventional Th1 cells to become cytotoxic CD4^+^ T cells during primary infection in the lung requires local antigen presentation and exposure to IL-15 (65).

Since the original description of cDC1 and the availability of mice that lack this subset, they have been prominently linked to cross presentation and the production of IL-12 (2,7,66,67). The studies presented here indicate that while cDC1 are not required for the initial priming of *Cryptosporidium*-specific CD4^+^ T cells they provide secondary signals required for CD4^+^ T cell expansion, gut homing, and expression of Th1 factors like T-bet and IFN-γ. Similarly, a recent report highlighted that *Irf8+32^-/-^* mice infected with *Cryptosporidium tyzzeri* develop parasite-specific CD4^+^ T cell responses but these are T-bet deficient (37). However, unlike the present study, these CD4^+^ T cell populations can access the SILP (37). This different requirement of cDC1 for homing and retention may reflect species differences (*C. parvum* is adapted to humans and cattle and *C. tyzzeri* causes persistent infection in mice) but for both species, cDC1s have a profound influence on the process that generate protective CD4^+^ T cell responses in the gut. Given the importance of cDC1-derived IL-12 in the development of Th1 responses (68), it was an initial surprise that CD4^+^ T cell responses in the absence of IL-12p40 did not phenocopy those from *Irf8+32^-/-^* mice. This difference underscores IL-12-independent functions for cDC1s to support CD4^+^ T cell responses in the gut. One plausible mechanism is the ability of gut-associated CD103^+^ cDC1s to produce retinoic acid that promotes expression of the gut homing integrins that drive T cell recruitment to the small intestine (9). In addition, the CD40-CD40L interaction is an important pathway for the control of *Cryptosporidium* (69), which has a prominent role in DC licensing (70) and in the mesLN can promote cDC1 IL-27 production (71). The observation that IL-27 is required shortly after weaning for the priming of Th1 cells and their accumulation in the SILP (8) suggest that this factor may also be relevant to the ability of cDC1 to promote Th1 responses to a mucosal pathogen.

Because *Cryptosporidium* is restricted to IECs and does not infect professional APCs, there are basic questions about how parasite-derived antigens gain access to the mLN and whether APCs at the site of infection can acquire parasite antigens and support local CD4^+^ T cell function (54). Interestingly, IECs can express MHCII, and differentiation of CD4^+^ T cells into CD4^+^CD8αα IELs within the gut requires MHCII expression on IECs (72). During *Cryptosporidium* infection, IFN-γ signaling on IECs leads to higher MHCII and CIITA expression (32,73 Preprint), which may facilitate the ability of IECs to engage effector CD4^+^ T cells within the IEL. Alternatively, during murine adenovirus (AdV) infection, viral specific effector Ly6A^+^ CD4^+^ T cells acquire “innate-like” functions in the gut and can produce IFN-γ after exposure to IL-12 plus IL-18 without TCR stimulation (52). It is notable that that while IEC-derived IL-18 is important for ILC production of IFN-γ and innate resistance to *Cryptosporidium* (28,32,74,75), the blockade of IL-18 did not result in reduced CD4^+^ T cell production of IFN-γ. Rather, IL-12 appears to be a dominant signal three required for these protective responses, but whether this depends on local TCR signals remains to be determined.

While the CD4^+^ T cell response to *Cryptosporidium* is characterized by a dominant Th1 phenotype, there is also the presence of a small population of Th17 cells, that is perhaps most apparent in the *Irf8+32^-/-^* mice infected with *C. tyzzeri* (37). Likewise, in the studies presented here a small percentage of SMARTA T cells were RORγT^+^ but there is little appreciation of the role of Th17 associated cytokines (IL-17, IL-22, GM-CSF) in resistance to *Cryptosporidium* (76,77). Nevertheless, this mixed Th1/Th17 response is reminiscent of recent work that compared the T cell responses in the gut to *Salmonella*, *Citrobacter* and *Nippostronglyus brasiliensis* which concluded that these diverse challenges can result in “mixed” responses that consist of cells that display Th1-like and Th17-like features (78). These observations led to the suggestion that pathogens exert broad influence on the state of CD4^+^ T cells that cannot be captured merely by classifying pathogens as eliciting “Th1” or “Th17” responses (78). It is possible that this may be a byproduct of a less differentiated state associated with mucosal priming (see earlier discussion) but we and others have noted the presence of a prominent IFN-γ-independent CD4^+^ T cell dependent mechanism to control *Cryptosporidium* (31,33). The basis for this phenomenon is unclear (73), but the availability of these transgenic parasites should facilitate studies to directly assess what IFN-γ-independent functions of CD4^+^ T cells mediate control of *Cryptosporidium*.

### Materials and methods Mice

C57BL/6 (stock no: 000664), Nur77-GFP reporter mice (stock no: 016617), CD45.1 C57BL/6 mice (stock no: 002014), SMARTA CD45.1 mice (stock no: 030450), *Irf8+32*^-/-^ mice (stock no: 032744), *Il12b*^/-^ mice (stock no: 002693), and *Ifng*^-/-^ (stock no: 002287) were purchased from Jackson Laboratories and the SMARTA CD45.1, SMARTA x IFN-g-CD90.1, SMARTA x Nur77-GFP CD45.1, *Irf8+32*^-/-^, *Il12b*^-/-^, and *Ifng*^-/-^ mice were maintained in-house. IFN-g-CD90.1 knock-in reporter mice were provided by Dr. Phillip Scott but originated in the laboratory of Dr. Casey Weaver (46). Mice used in this study were males or females ranging from 4 to 10 weeks. All mice were age and sex matched within individual experiments. All protocols for animal care were approved by the Institutional Animal Care and Use Committee of the University of Pennsylvania (protocol #805405 and #806292).

### Plasmid construction

To see full list of primers used for plasmid construction, see Supplementary Table 1. To generate *Cp*-2W1S and *Cp*-gp61: the pLIC_SIINFEKL_gp61_HA mNeon and pLIC_HA_2W1S_SIINFEKL plasmids, pLIC_HA_t2A_mNeon (79) was amplified using primers 1 and 2 containing the SIINFEKL_gp61 sequence or primers 3 and 4, containing the 2W1S_SIINFEKL sequence, respectively. PCR fragments were subsequently re-ligated by T4 DNA ligase. Repair templates were amplified using primers 5 and 6 with 30 bp overhangs either side of a Cas9 guide-induced double strand break in the MEDLE2 locus (Cgd5_4590). The MEDLE2 targeting guide plasmid was used as previously described (42).

To generate ma*Cp*-ova-gp61: the repair template encodes the last 113 bp of the pheRS gene (cgd_3320, recodonized) (49 Preprint) including the mutation that confers resistance (CTT to GTT at nucleotide position 1444; corresponding to L482V) to BRD7929. This short sequence is followed by MEDLE2-ova-gp61-HA (where ova is SIINFEKL and gp61 is GLKGPDIYKGVYQFKSVEFD) driven by the *C. parvum enolase* promotor, which was inserted using Gibson assembly into the pheRS^R^ plasmid previously generated (49). The repair template was amplified using primers 7 and 8 with 30 bp overhangs on either side of a Cas9 guide-induced double strand break in the pheRS locus. The pheRS targeting guide plasmid was used as previously described (49 Preprint).

### Isolation of transgenic parasites

Transgenic parasites were derived as previously described (55). For *Cp-*2W1S and *Cp-*gp61, 5×10^7^ *C. parvum* oocysts (Bunchgrass, USA) were incubated in 1:4 ice cold bleach in PBS on ice for 10 minutes, washed in 1x PBS and incubated in 0.8% sodium taurodeoxycholate to excyst sporozoites. For ma*Cp*-ova-gp61, sporozoites were excysted by washing 60×10^7^ mCherry mouse-adapted *C. parvum* (ma*Cp,* described previously (32)) in 1x PBS and then incubating in 10 mM HCl for 45 minutes at 37℃. Next, oocysts were pelleted, washed twice in 1x PBS, and incubated in 0.2 mM sodium taurodeoxycholate and 20 mM sodium bicarbonate in 1X PBS for 1 hour at 37℃ to excyst ma*Cp* sporozoites. Excysted sporozoites were resuspended in transfection buffer supplemented with a total of 100 μg DNA (50 μg of Cas9/gRNA plasmid and 50 μg of repair template generated by PCR) and electroporated using an Amaxa 4D nucleofector (Lonza, Basel, Switzerland). *Cp-*2W1S and *Cp-*gp61 parasites carrying a stable transgene were selected with paromomycin (16 g/L) added to the drinking water of orally-infected *Ifng*^-/-^ mice. ma*Cp*-ova-gp61 were selected using BRD7929 (48) orally gavaged daily (10 mg/kg/day) for the first six days after transfection. Oocysts were purified from feces using sucrose floatation followed by cesium chloride gradient as described previously (31).

### Mouse infection and measurement of parasite burden

For experiments using *Cp-*2W1S or *Cp-*gp61, *Ifng*^-/-^ mice were infected with 1×10^4^ oocysts by oral gavage. For experiments using ma*Cp*-ova-gp61, mice were infected with 5×10^4^ oocysts by oral gavage. To quantify fecal oocyst shedding, 20 mg of pooled feces was suspended in 1mL lysis buffer. Samples were shaken with glass beads for 5min, then combined 1:1 ratio with Nano-Glo Luciferase solution (Promega). A ProMega GloMax plate reader was used to measure luminescence. Pooled samples from cages were used because previous studies have demonstrated mice within each cage are equally infected (80).

### T cell transfers

For T cell transfers, SMARTA CD45.1 mice, SMARTA CD45.1 interbred with CD45.1/Nur77-GFP reporter mice (where indicated), or SMARTA CD45.1 interbred with IFN-g-CD90.1 (where indicated) were used. To purify SMARTA T cells, spleens and lymph nodes were isolated by dissociation over a 70 μm filter. Red blood cells were lysed by incubation for 5 minutes at room temperature in 1mL of ACK lysis buffer (0.864% ammonium chloride (Sigma-Aldrich) diluted in sterile-deionized H2O) and then washed in complete RPMI (cRPMI, 10% fetal calf serum, 0.1% beta-2-mercaptoethanol, 1% non-essential amino acids, 1% sodium pyruvate, and 1% penicillin-streptomycin). SMARTA T cells were enriched by magnetic activated cell sorting (MACS) using the EasySep^TM^ Mouse CD4+ T cell Isolation Kit (Stem Cell technologies). SMARTA purity was verified (>80%) using flow cytometry for TCR Vα2 and Vβ8.3 expression. 2×10^4^-1×10^6^ were transferred by intravenous injection into recipient mice. If using CellTrace Violet, SMARTA T cells were fluorescently labelled using the CTV labeling kit (Thermo Fisher Scientific) prior to i.v. transfer.

### Tissue isolation and flow cytometry

To harvest the epithelial/IEL fraction, single-cell suspensions were prepared from ileal sections by shaking diced tissue at 37°C for 20 minutes in Hank’s Balanced Salt Solution (HBSS) with 5 mM EDTA and 1 mM DTT. Cell pellets were then passed through 70 μm and 40 μm filters. For lamina propria harvesting, the remaining intestine after epithelial layer isolation was washed in HBSS plus 10mM HEPES to remove EDTA, minced with scissors, and incubated in complete RPMI with 0.16 mg/mL Liberase TL (Roche) and 0.1 mg/mL DNase I (Roche) with shaking for 30 minutes at 37°C. Cell pellets were then passed through 70 μm and 40 μm filters. Ileal draining mesenteric lymph nodes and Peyer’s patches from the ileum as well as spleens were harvested and dissociated through 70 um filters, then washed with cRPMI. Cells were washed in FACS buffer (1x PBS, 0.2% bovine serum albumin, 1 mM EDTA), and incubated in Fc block (99.5% FACS Buffer, 0.5% normal rat IgG, 1 µg/ml 2.4G2) at 4°C for 15 minutes prior to staining. Cells were stained for cell death using GhostDye Violet 510 Viability Dye or GhostDye Red 780 Viability Dye (TONBO Biosciences) in 1x PBS at 4°C for 15 minutes. Cells were washed after cell death staining and surface antibodies were added and stained at 4°C for 30 minutes. If tetramer staining, prior to Fc block cells were washed with PBS+2% FCS and then incubated in RPMI+10% FCS with tetramer at 37°C for 45 minutes. If intracellular staining for transcription factors or cytokines was performed, after surface staining cells were fixed using the eBioscience Foxp3 Transcription Factor Fixation/Permeabilization Concentrate and Diluent (ThermoFisher Scientific) for 45 minutes at 4°C. Cells were then stained for transcription factors in 1x eBioscience Permeabilization Buffer (ThermoFisher Scientific) at room temperature for 30 minutes. Cells were then washed in Permeabilization Buffer and then washed in FACS buffer prior to acquisition. For intracellular cytokine staining of T cells, prior to plating for staining, single cell suspensions from each tissue were plated in a 96-well plate and incubated with gp61 peptide and Protein Transport Inhibitor Cocktail (eBioscience) in cRPMI for 3.5 hours at 37°C. For intracellular cytokine staining of DCs, prior to plating for staining, single cell suspensions from each tissue were plated in a 96-well plate and incubated with Brefeldin A (Sigma Aldrich) in cRPMI for 6 hours at 37°C.

Cells were stained using the following fluorochrome-conjugated antibodies: ef450 CD45.1 (clone 104, Invitrogen), BV605 CD44 (clone IM7, eBioscience), BV711 CD45.1 (clone A20, Biolegend), Af700 CD90.2 (clone 3-H12, Biolegend), PE CD8a (clone 53-6.7, eBioscience), BUV395 NK1.1 (clone PK136, BD), BUV395 CD19 (clone ID3, BD), BUV805 CD (clone GK1.5, BD), ef450 IL-18Ra (clone P3TUNYA, eBioscience), BV785 CD90.2 (clone 3-H12, Biolegend), APC-e780 CD90.1 (clone H1S51, eBioscience), BUV395 EpCAM (clone G8.8, BD), PE IL-17A (clone TC11-18H10.1, BD), BV711 TNFa (clone MP6-XT22, Biolegend), PE CD45.1 (clone A20, BD), PE-Cy5 T-bet (clone 4B10, Invitrogen), PE-Cy5.5 Foxp3 (clone FJK-16.S, eBioscience), PE-Cy7 IL-2 (JES6-5H4, eBioscience), BUV496 CD4 (clone GK1.5, BD), BUV737 IFN-γ (XMG1.2, BD), BUV805 CD8a (53-6.7, BD), FITC CD4 (clone GK1.5, eBioscience), biotin CXCR5 (clone SPRCL5, eBioscience), streptavidin BV421 (Biolegend), BV786 RORγT (clone Q31-378, BD), Af647 GATA3 (clone L50-823, BD), PE Bcl-6 (clone 1G19E/A8, Biolegend), PE-Cy7 PD-1 (clone J43, eBioscience), BUV496 CD44 (clone IM7, BD), FITC Ly6C (clone AL-21, BD), ef450 CD11b (clone M1/70, eBioscience), BV605 CD103 (clone 2E7, Biolegend), BV650 XCR1 (clone ZET, Biolegend), BV711 CD26 (clone H194-112, BD), APC IL-12p40/p70 (clone C15.6, BD), I-A/E Af700 (clone M5/114.15.2, eBioscience), APC-Cy7 SIRPa (clone P84, Biolegend), PE CD40 (clone 1C10, eBioscience), PE-Cy5 CD86 (clone GL1, Biolegend), PE-Cy7 CD64 (clone X54-5/7.1, Biolegend), BUV395 CD3e (clone 145-2C11, BD), BUV395 B220 (clone RA3-6B2, BD), BUV737 CD11c (clone HL3, BD), Af488 CD25 (clone PC61.5, eBioscience), APC CD45.2 (clone 104, eBioscience), APC-Cy7 CD8a (clone 53-6.7, Biolegend), PE RORγT (clone B2D, eBioscience), PE-Cy7 CD27 (clone LG.7F9, eBioscience), BUV737 CD69 (clone H1.2F3, BD), FITC CD8a (clone 53-6.7, eBioscience), BV421 IL-23R (clone 12B2B64, Biolegend), BV785 CD25 (clone PC61.5, Biolegend), APC-Cy7 CD44 (clone IM7, Biolegend), PE LPAM-1 (clone DATK32, eBioscience), PE-Cy7 IL-18Ra (clone P3TUNYA, eBioscience), PE CD40L (clone MR1, Biolegend), PE-Cy5 CD44 (clone IM7, BD), PE-Cy7 CD69 (clone H1.2F3, eBioscience), PerCP-ef710 LPAM-1 (clone DATK32, eBioscience), APC TCR Vβ5.1 (clone MR9-4, Biolegend), PE TCR Vβ8.3 (clone 1B3.3, Biolegend), PE-Cy7 CD90.1 (clone H1S51, eBioscience), BUV563 CD8a (clone 53-6.7, BD), Af488 IL-12RB2 (clone 305719, R&D Systems), PerCP-ef710 IL-18Ra (clone P3TUNYA, eBioscience), APC IL-21R (clone 4A9, Biolegend), BV421 XCR1 (clone ZET, Biolegend), APC SIRPa (clone P84, Biolegend), APC-e780 CD11c (clone N418, eBioscience), BV421 SLAM (clone TCF-12F12.2, Biolegend), APC CD40L (clone MR1, Invitrogen), Af700 Ly6A/E (clone D7, eBioscience), APC-ef780 TCRβ (clone H57-597, eBioscience), PE-Cy7 IL-18Ra (clone P3TUNYA, eBioscience), BUV805 CD8a (clone 54-6.7, BD) BUV496 CD44 (clone IM7, eBioscience). Endogenous 2W1S:I-Ab responses were measured by 2W1S:I-Ab tetramer conjugated to PE or APC (NIH Tetramer Core). Data were collected on a FACSCanto, LSRFortessa, FACSymphony A3 Lite, or FACSymphony A5 (BD Biosciences) and analyzed with FlowJo v10 software (TreeStar). CD4^+^ T cells (CD4^+^TCRb^+^EpCAM^-^NK1.1^-^CD8α^-^CD19^-^) were downsampled to 1000 cells using Downsample plugin in FlowJo prior to UMAP. UMAP and X-Shift plugins were utilized in FlowJo for dimensionality reduction and unsupervised clustering, and X-Shift was visualized by Cluster Explorer.

### Immunofluorescence imaging

Coverslips seeded with human ileocecal adenocarcinoma cells (HCT-8) (ATCC) were infected when 60-80% confluent with 300,000 purified oocysts (bleached, washed, and resuspended in DMEM medium (Gibco) supplemented with 10% CCS (Cytiva). At 24 hours, cells were washed with PBS, and successively fixed with 4% paraformaldehyde (Electron Cytometry Sciences) for 15 minutes and permeabilized with 0.1% Triton X-100 (Sigma) in PBS for 10 minutes. Coverslips were blocked with 4% bovine serum albumin (BSA) (Sigma) in PBS. Antibodies were diluted in 1% BSA in PBS. Rat monoclonal anti-HA (MilliporeSigma) was used as primary antibody (1:1000) and goat anti-rat polyclonal Alexa Fluor 594 (Thermo Fisher) as secondary. Host and parasite nuclei were stained with Hoechst 33342 (Thermo Fisher). Slides were imaged using a Leica DM6000B widefield microscope.

### Statistics

Statistical significance was calculated using unpaired t test with Welch’s correction for comparing groups of 2 or ANOVA followed by multiple comparisons for comparing groups of 3 or more. Analyses were performed using GraphPad Prism v9.

## Supplemental material

Supplementary Table S1. shows oligonucleotides used for generating transgenic parasites used in this study. Figure S1 shows the integration PCR for *Cp*-2W1S and *Cp*-gp61 transgenic parasites. Figure S2 shows representative flow cytometry plots from CD90.1-*Ifng* reporters (SFig 2A), from peptide-stimulated mesLN SMARTA T cells (SFig 2B), and a summary of TNF-α production by peptide-stimulated SMARTA T cells from the SILP and mesLN. Figure S3 shows a construct map for generating transgenic ma*Cp*-ova-gp61 from ma*Cp* and the integration PCR for this strain in SFig 3A-B. SFig 3C-E relates to the UMAP experiments shown in Figure 3. Figure S4 shows additional data related to Figure 4. Figure S5 is related to Figure 5, showing an absence of IL-12p40 expression in cDC2s and modest expression in macrophages. Figure S6 is related to Figure 6 showing that *Irf8+32^-/-^* mice remain deficient during *Cryptosporidium* infection. Figure S7 is related to Figure 7 showing intact gut-homing but impaired maintenance of SMARTA T cells in *Il12b^-/-^* mice, and the impact of IL-18 and/or IL-12p40 blockade on IL-18Ra expression in SMARTA T cells.

## Acknowledgements

This work was supported in part by the National Institutes of Health with grants to C.A.H. and B.S. (U01AI163671 and R01AI148249), to B.S. (R01AI112427), to C.A.H. (U01AI160664, R01AI157247), a fellowship to I.S.C. (F30AI169744), training grant support to B.E.H. (T32AI007532), J.A.G. (T32AI055400), K.M.O (T32AI055428), and M.I.M. (T32AI007632), fellowships from the Swiss National Science Foundation to S.S. (P2BEP3_191774 and P500PB_211097), and a fellowship from the Canadian Institutes of Health Research (MFE-176621) and a Postdoctoral Training award from the Fonds de Recherche du Québec – Santé (300355) to R.D.P. B.S. and C.A.H. are supported by the Commonwealth of Pennsylvania.

## Author Contributions

I.S.C., B.S., and C.A.H. conceived of these studies. I.S.C. conducted experiments with help from B.A.W., B.E.H., K.M.O., R.D.P., and M.I.M. B.A.W., I.S.C., and S.S. engineered transgenic parasites. I.S.C., B.E.H., J.A.G., and D.A.C. bred, maintained, and provided mice for these studies. All authors approved the final manuscript and agree to be accountable for all aspects of the work in ensuring that questions related to the accuracy or integrity of any part of the work are appropriately investigated and resolved.

---

**Supplementary Table S1:**
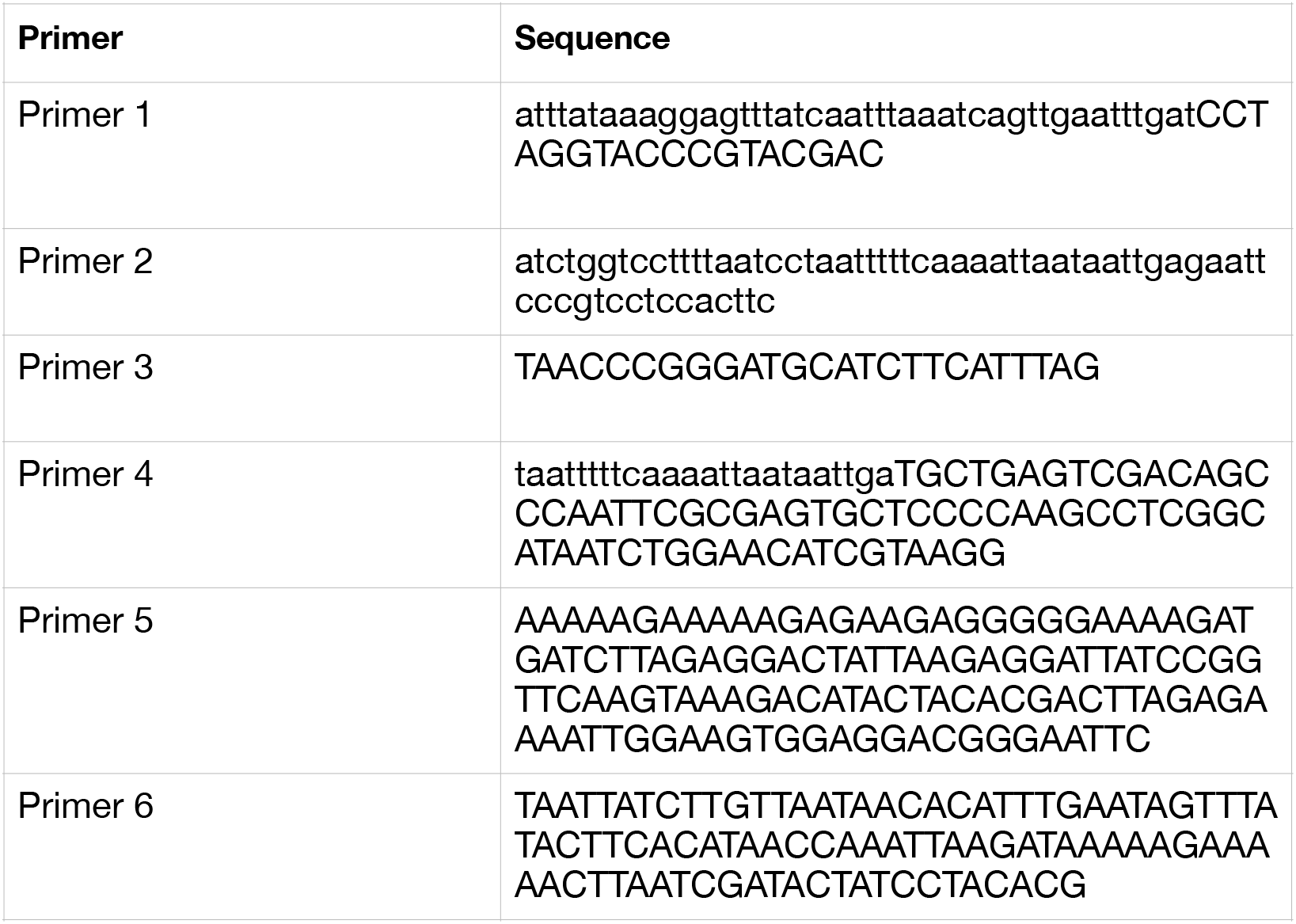
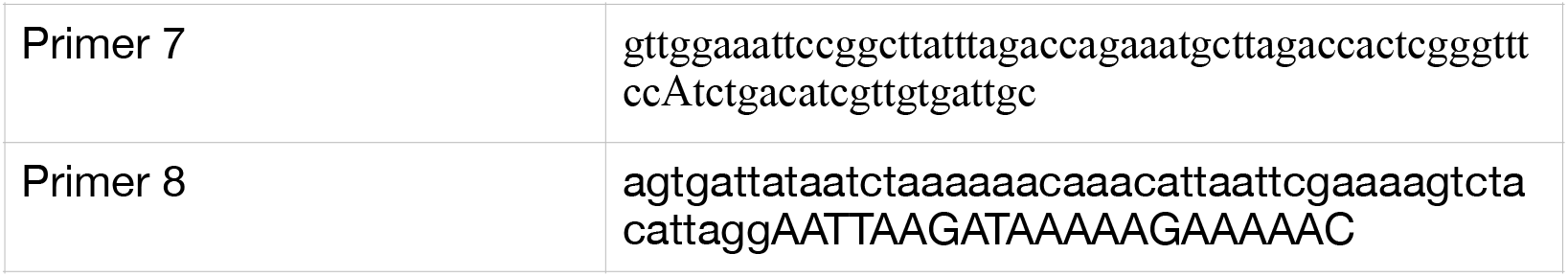
Oligonucleotides for guide RNAs and primers for repair template amplification used in this study.

## Supplementary Figure

**Supplementary Figure 1.**
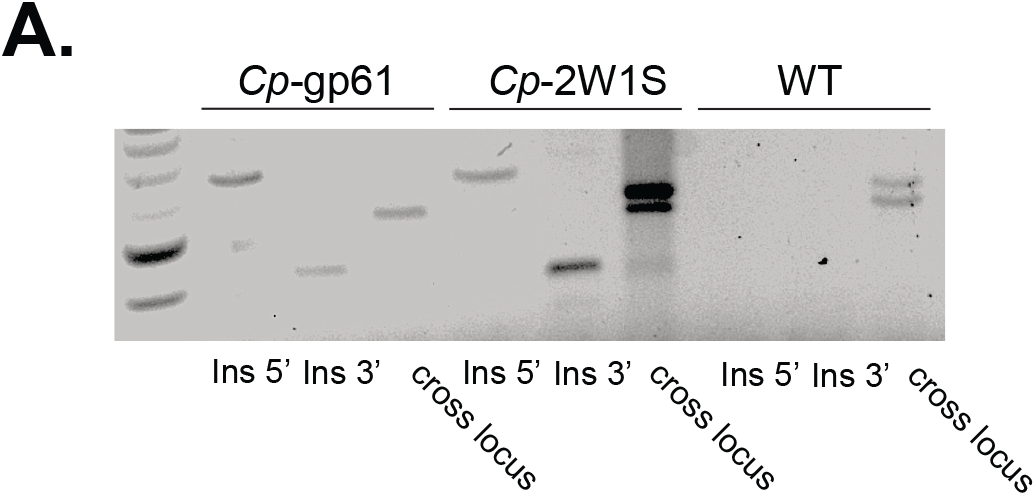
**A.** Integration PCR gel. PCR mapping using genomic DNA from wild-type (WT) and transgenic parasites (*Cp*-gp61 and *Cp*-2W1S) showing diagnostic amplicons of the insertion locus. Primer binding sites and expected amplicon sizes are indicated in Figure 1A on the genetic maps. Because there are multiple copies of MEDLE2 in the *C. parvum* genome, loss of the WT cross-locus band in transgenic parasites is not necessarily expected.

**Supplementary Figure 2.**
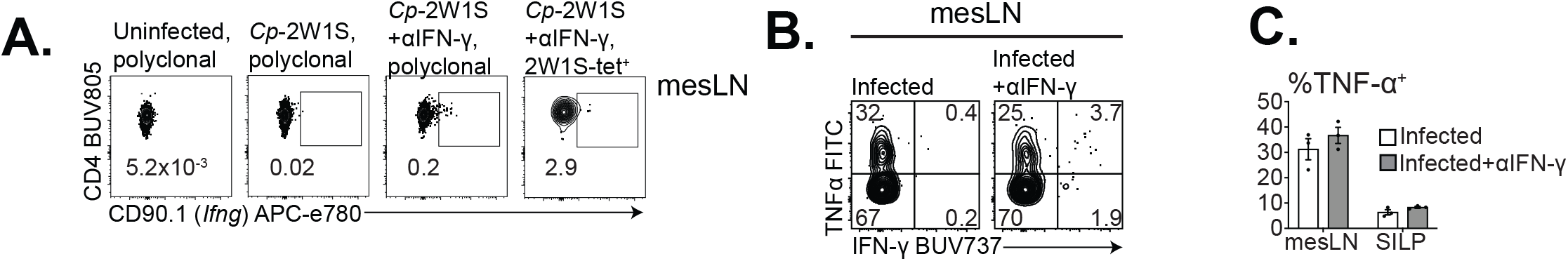
**A.** Representative flow plots showing CD90.1 expression in polyclonal CD4^+^ T cells (first two columns) or 2W1S:I-Ab tetramer^+^ cells (last column) in the mesLN of uninfected mice (first column), infected mice untreated (second column) or infected mice treated with αIFN-γ (last two columns). Gating: Singlets, Live^+^, CD19^-^, NK1.1^-^, EpCAM^-^, CD90.2^+^, CD8a^-^, CD4^+^, 2W1S:I-Ab^+^. **B-C.** WT B6 mice received 2×10^4^ CD45.1^+^ SMARTA T cells 1 day prior to infection with 5×10^4^ ma*Cp*-ova-gp61 and were left untreated or treated with 1mg/mouse of αIFN-γ 1 day prior to infection and 2, 5, and 8 dpi. At 10 dpi mesLN and SILP were harvested for flow cytometry and cells were stimulated with exogenous gp61 peptide for 3 hours followed by intracellular cytokine staining and flow cytometry. **B.** Representative flow plots from the mesLN. **C.** Summary showing the percentage of SMARTA cells from the SILP or mesLN staining TNFα^+^ after peptide stimulation. Gating: Singlets, Live^+^, CD19^-^, NK1.1^-^, EpCAM^-^, CD90.2^+^, CD8a^-^, CD4^+^, CD45.1^+^. Statistical significance was determined by two-way ANOVA and multiple comparisons. p≤0.05, ** p≤0.01, *** p≤0.001, **** p≤0.0001.

**Supplementary Figure 3.**
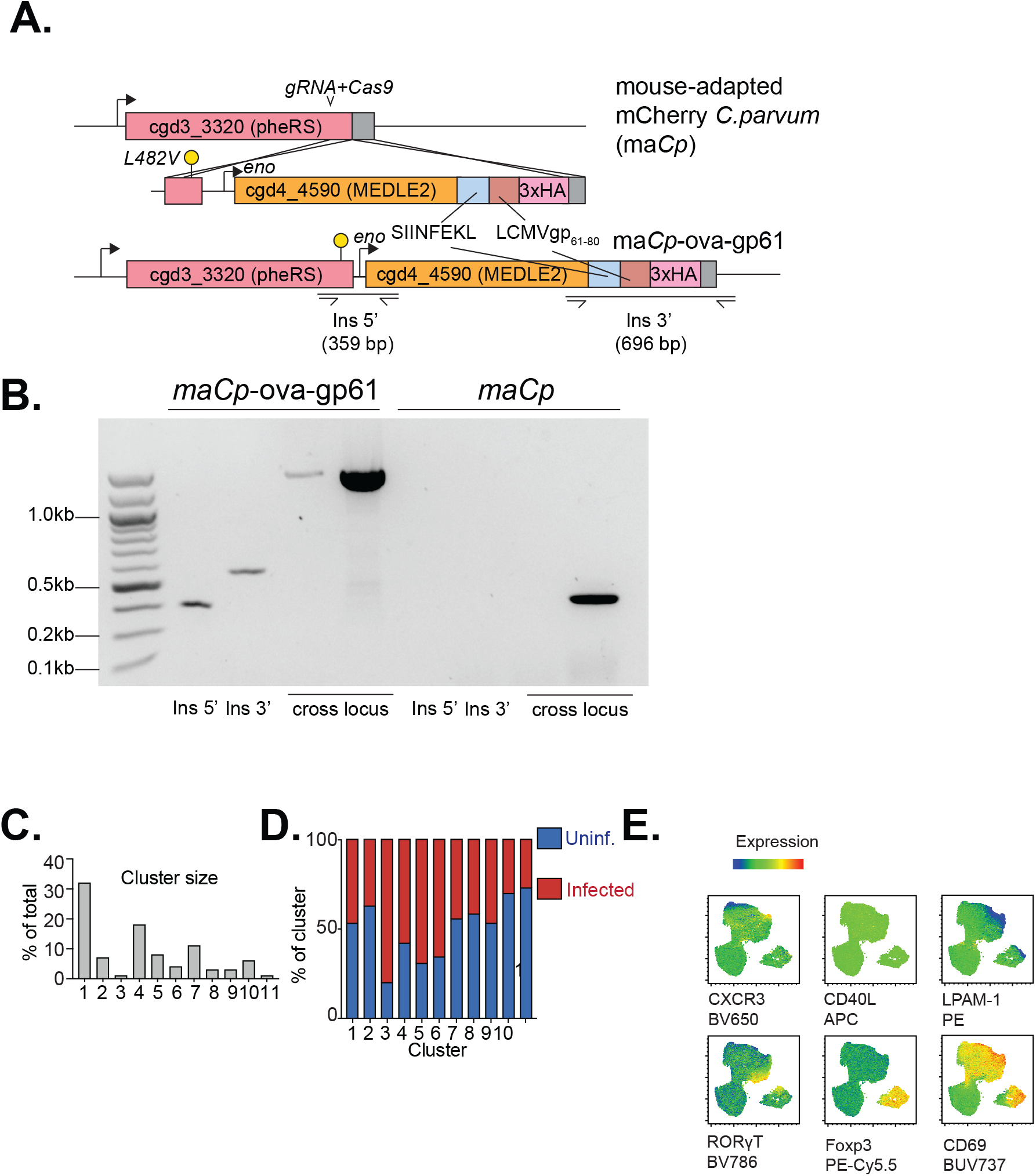
**A.** Genetic construct of transgenic ma*Cp*-ova-gp61 engineered to alter the pheRS gene L at position 482 to V to confer resistance to BRD7929. This construct included the full MEDLE2 gene tagged with SIINFEKL-gp61-3xHA. **B.** Integration PCR gel. PCR mapping using genomic DNA from ma*Cp* and transgenic parasites (ma*Cp*-ova-gp61) showing diagnostic amplicons of the insertion locus. Primer binding sites and expected amplicon sizes are indicated in SFig 1A on the genetic maps. **C.** Percentage of total CD4^+^ T cells represented by each cluster from UMAP experiments described in Figure 3. **D.** Percentage of each cluster represented by CD4^+^ T cells from uninfected (blue) or infected (red) samples. **E.** UMAP from Figure 3A colored by expression levels of each marker listed beneath each panel.

**Supplementary Figure 4.**
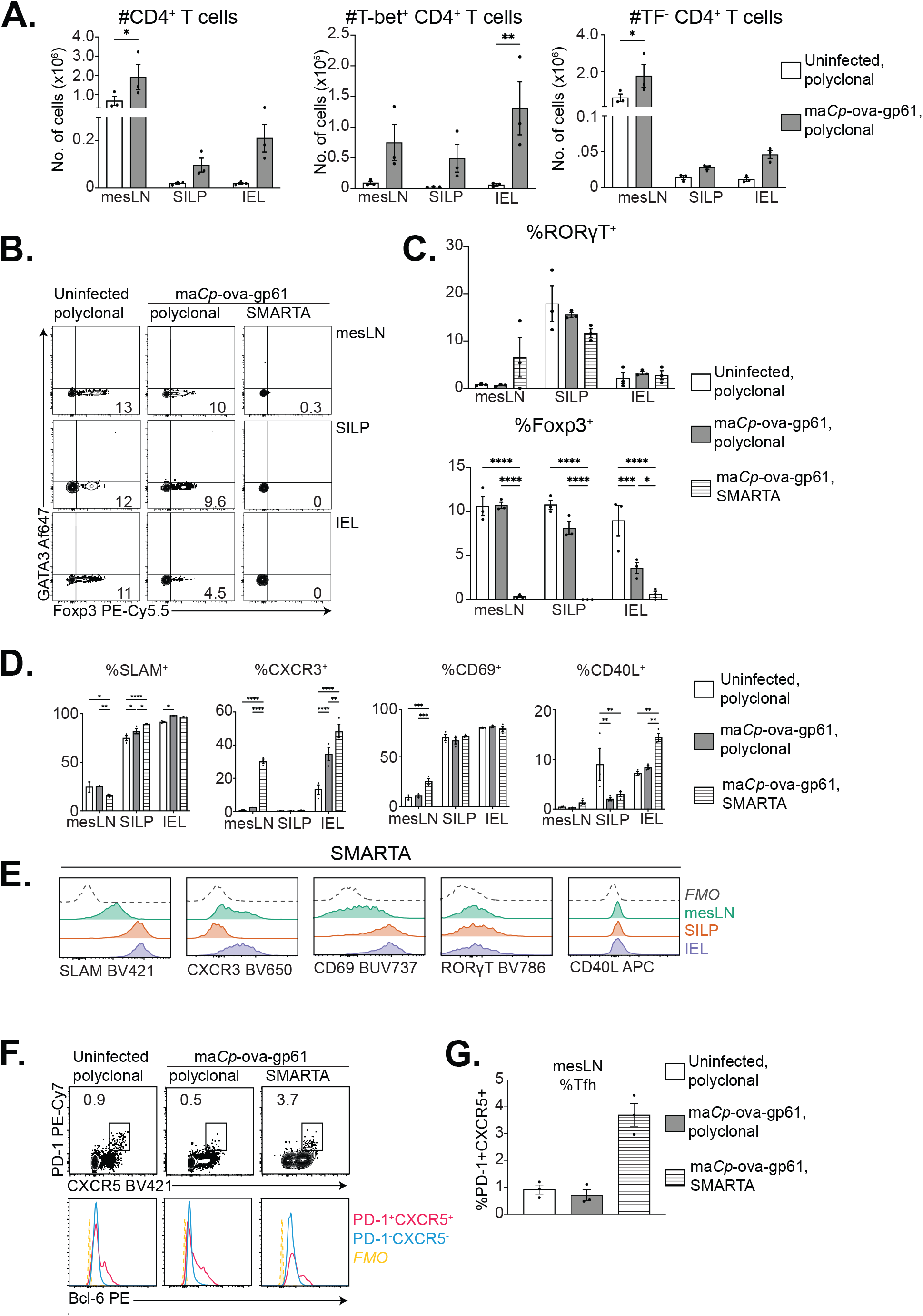
**A.** Numbers of total, T-bet^+^, or TF^-^ (T-bet^-^RORγT^-^ GATA3^-^Foxp3^-^) CD4^+^ T cells in uninfected or infected mice at 10 dpi with 5×10^4^ ma*Cp-*ova-gp61. **B.** Representative flow plots of Foxp3 vs. GATA3 in polyclonal CD4^+^ T cells from uninfected mice, polyclonal CD4^+^ T cells from infected mice, or SMARTA T cells (CD45.1^+^) from infected mice organized by tissue (rows). Gating: Singlets, Live^+^, CD19^-^, NK1.1^-^, EpCAM^-^, CD90.2^+^, CD8a^-^, CD4^+^, CD45.1^+.^. **C.** Percentage of RORγT^+^ or Foxp3^+^ in polyclonal CD4^+^ T cells from uninfected mice (white), polyclonal CD4^+^ T cells from infected mice (gray), or SMARTA T cells (horizontal lines). **D.** Percentage of cells positive for SLAM, CXCR3, CD69, or CD40L of polyclonal CD4^+^ T cells from uninfected mice (white), infected mice (gray), or SMARTA T cells (stripes). **E.** Histograms of protein expression by flow cytometry among SMARTA T cells from the mesLN (teal), SILP (orange) or IEL (purple). Gating: Singlets, Live+, CD8a^-^EpCAM^-^NK1.1^-^ CD19^-^CD4^+^TCRβ^+^CD45.1^+^. **F-G**. Representative flow cytometry plots of polyclonal CD4^+^ T cells or SMARTA T cells staining for Tfh markers PD-1, CXCR5, and Bcl6 with summary in (G). Gating: Singlets, Live^+^, CD19^-^, NK1.1^-^, EpCAM^-^, CD90.2^+^, CD8a^-^, CD4^+^, CD45.1^+.^. For A-C, F-G, representative of 2 independent experiments; for D-E, one independent experiment. n=3 mice/ group. Statistical significance was determined by two-way ANOVA and multiple comparisons. p≤0.05, ** p≤0.01, *** p≤0.001, **** p≤0.0001. Statistical significance was determined by two-way ANOVA and multiple comparisons. p≤0.05, ** p≤0.01, *** p≤0.001, **** p≤0.0001.

**Supplementary Figure 5.**
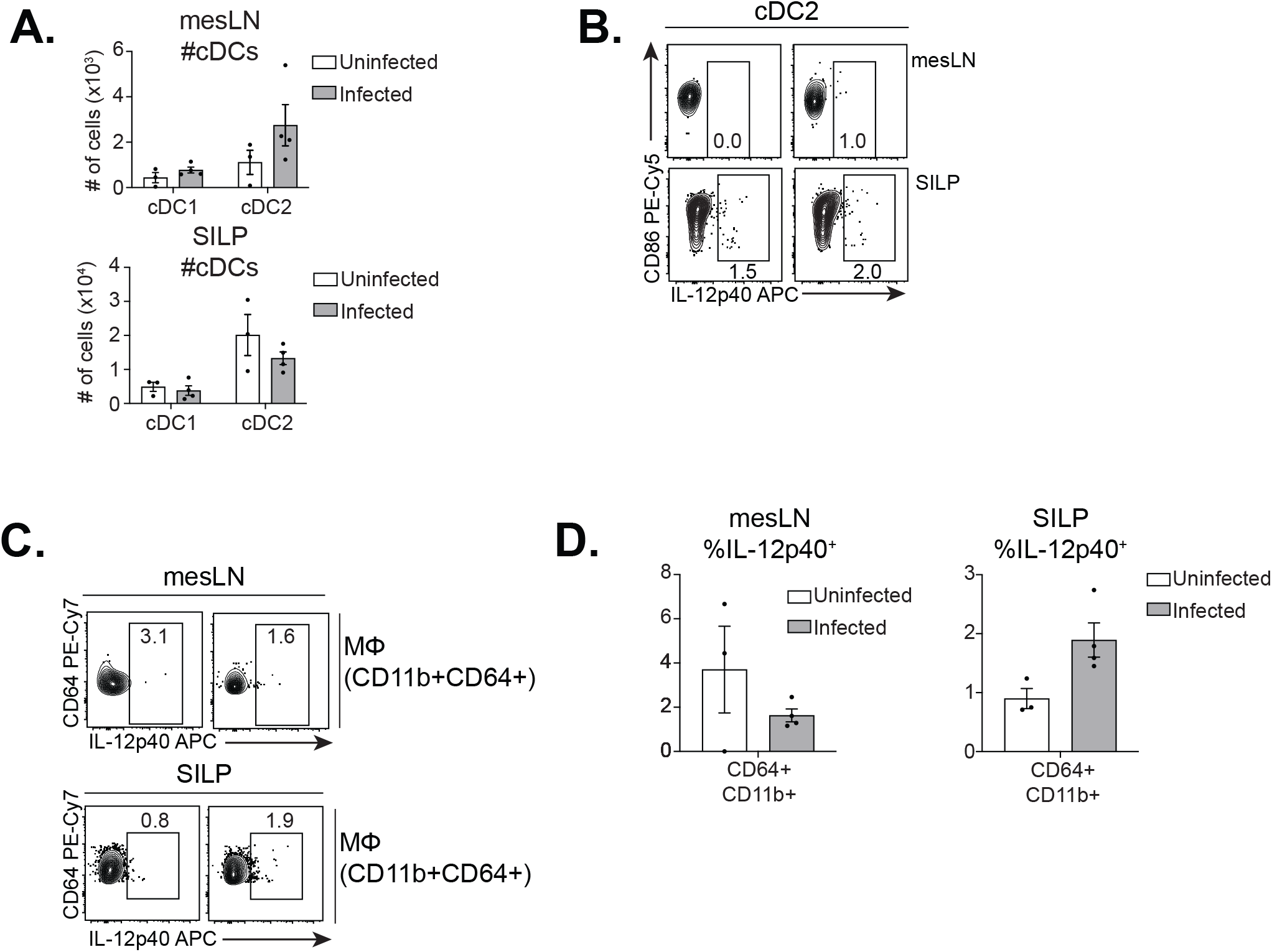
**A-D.** WT B6 mice were left uninfected or infected with 5×10^5^ ma*Cp* oocysts. At 4 dpi mesLN and SILP cells were isolated, plated with brefeldin A (BFA) for 6 hours, and analyzed by flow cytometry for surface markers and IL-12p40 expression. **A.** Total numbers of cDC1s or cDC2s in the mesLN (top) and SILP (bottom). **B.** IL-12p40 flow plots from the mesLN and SILP in cDC2s. Gating for A-B: Singlets, Live+, CD3e^-^, NK1.1^-^, EpCAM^-^, B220^-^, CD19^-^, CD90.2^-^, CD64^-^, MHCII-hi, CD11c^+^, CD26^+^, Ly6C^-^, XCR1^+^ (cDC1s) or SIRPα^+^ (cDC2s). **C.** IL-12p40 flow plots from the mesLN and SILP in macrophages. **D**. Summary of (C). Gating for C-D: Singlets, Live+, CD3e^-^, NK1.1^-^, EpCAM^-^, B220^-^, CD19^-^, CD90.2^-^, CD64^+^, CD11b^+^. Gating is based on FMOs taken from SILP for all colors except IL-12p40, for which positivity is based on samples from each tissue plated without BFA. Statistical significance was determined by two-way ANOVA and multiple comparisons. p≤0.05, ** p≤0.01, *** p≤0.001, **** p≤0.0001.

**Supplementary Figure 6.**
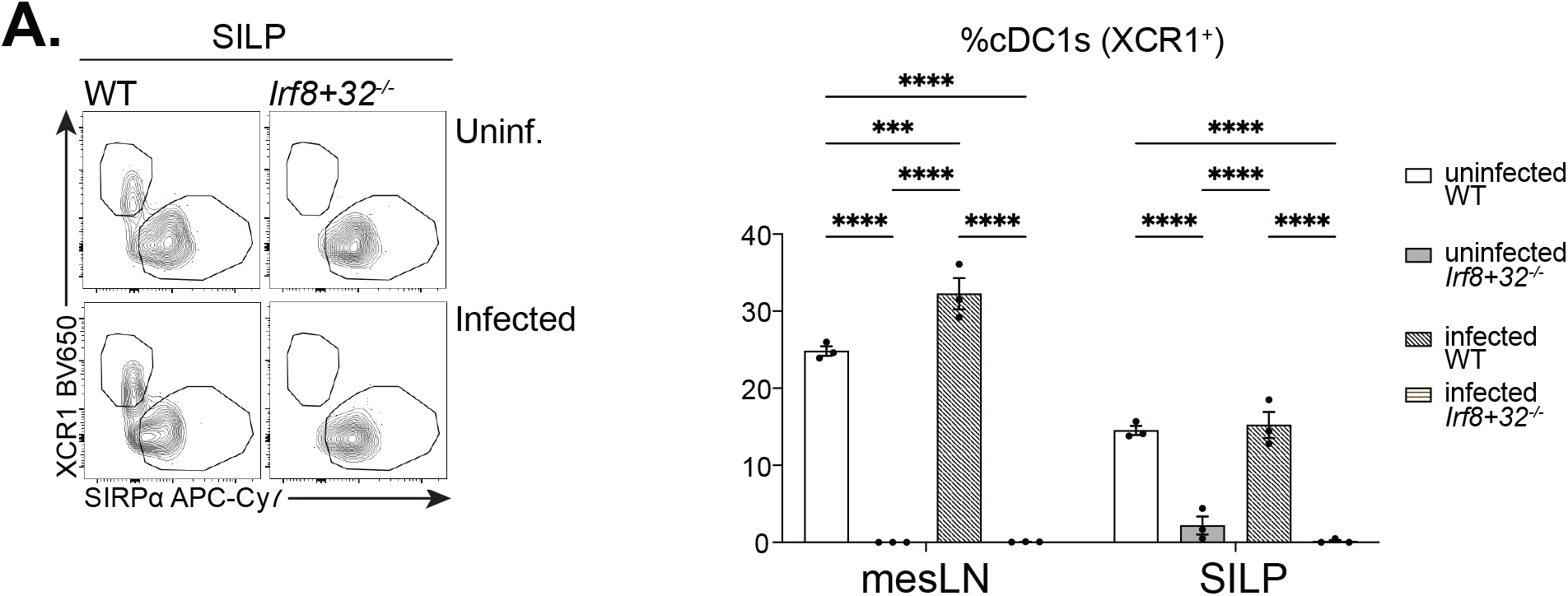
**A.** WT or *Irf8+32*^-/-^ mice were left uninfected or infected with 5×10^4^ ma*Cp* oocysts. At 3 dpi mice were sacrificed and percentage of cDC1s and cDC2s were quantified to confirm that cDC1s remained deficient in *Irf8+32*^-/-^ mice. Gating: Singlets, Live+, CD3e^-^, NK1.1^-^, EpCAM^-^, B220^-^, CD19^-^, CD90.2^-^, CD64^-^, MHCII-hi, CD11c^+^, CD26^+^, Ly6C^-^, XCR1^+^ (cDC1s) or SIRPα^+^ (cDC2s). Statistical significance was determined by two-way ANOVA and multiple comparisons. p≤0.05, ** p≤0.01, *** p≤0.001, **** p≤0.0001.

**Supplementary Figure 7.**
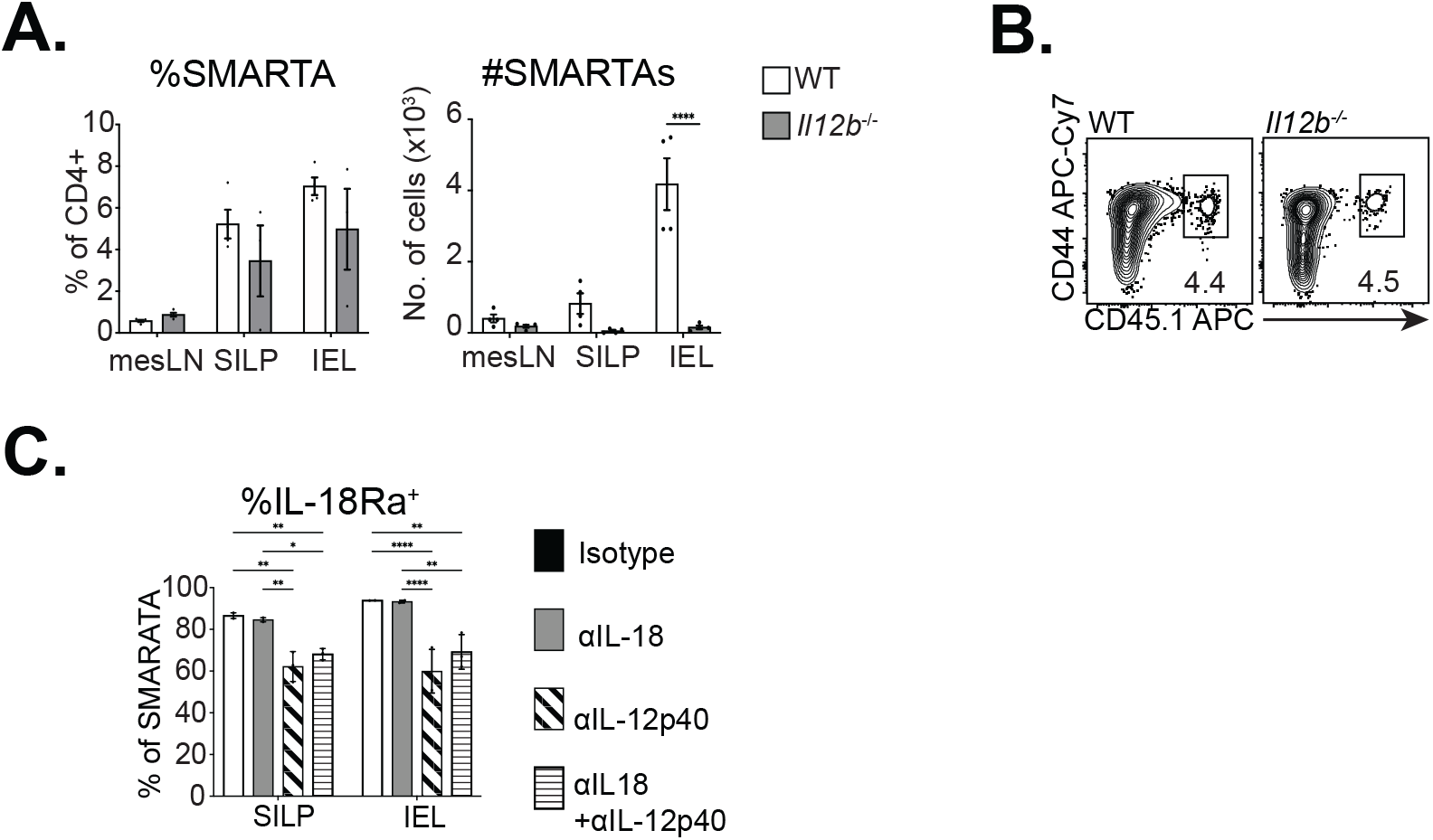
**A-B.** 1 day prior to infection with 5×10^4^ ma*Cp*-ova-gp61 oocysts, 2×10^4^ CD45.1+ SMARTA T cells were transferred into WT or *Il12b^-/-^* mice. At 10 dpi, mesLN, SILP, and IEL were isolated and analyzed by flow cytometry. Percentage of SMARTA T cells among CD4^+^ T cells and absolute numbers of SMARTA T cells from each genotype with representative flow plots from the SILP in (B). **C.** 1 day prior to infection with 10^4^ *Cp*-gp61, 5×10^4^ CD45.1^+^ SMARTA T cells that expressed CD90.1 as a reporter for *Ifng* expression were transferred into *Ifng*^-/-^ mice that were treated with isotype control (rat IgG2a), anti-IL-12p40, anti-IL-18, or anti-IL12p40+anti-IL-18 (1mg/mouse on d-1, or 2, 5, 8 dpi). Mice were also treated with 1mg/mouse anti-IFN-γ on 3 and 7 dpi. At 10 dpi, mesLN, SILP, and IEL were isolated and analyzed by flow cytometry. Shown is percentage of SMARTA T cells staining for IL-18Ra in each treatment condition. Gating: Singlets, Live+, NK1.1^-^, CD19^-^, EpCAM^-^, CD90.2^+^, CD8a^-^, CD4^+^, CD44-hi, CD45.1^+^. Statistical significance was determined by two-way ANOVA and multiple comparisons. p≤0.05, ** p≤0.01, *** p≤0.001, **** p≤0.0001.

## References

1. Yin X, Chen S, Eisenbarth SC. Dendritic Cell Regulation of T Helper Cells. Annu Rev Immunol. Mar 12 2021;doi:10.1146/annurev-immunol-101819-025146

2. Dudziak D, Kamphorst AO, Heidkamp GF, et al. Differential antigen processing by dendritic cell subsets in vivo. Science. Jan 5 2007;315(5808):107–11. doi:10.1126/science.1136080

3. Lehmann CHK, Baranska A, Heidkamp GF, et al. DC subset-specific induction of T cell responses upon antigen uptake via Fcgamma receptors in vivo. J Exp Med. May 1 2017;214(5):1509–1528. doi:10.1084/jem.20160951

4. Ferris ST, Durai V, Wu R, et al. cDC1 prime and are licensed by CD4(+) T cells to induce anti-tumour immunity. Nature. Aug 2020;584(7822):624–629. doi:10.1038/s41586-020-2611-3

5. Valdez Y, Mah W, Winslow MM, Xu L, Ling P, Townsend SE. Major histocompatibility complex class II presentation of cell-associated antigen is mediated by CD8alpha+ dendritic cells in vivo. J Exp Med. Mar 18 2002;195(6):683–94. doi:10.1084/jem.20010898

6. Theisen DJ, Davidson JTt, Briseno CG, et al. WDFY4 is required for cross-presentation in response to viral and tumor antigens. Science. Nov 9 2018; 362 (6415): 694–699. doi: 10.1126/science.aat5030

7. Mashayekhi M, Sandau MM, Dunay IR, et al. CD8alpha(+) dendritic cells are the critical source of interleukin-12 that controls acute infection by Toxoplasma gondii tachyzoites. Immunity. Aug 26 2011;35(2):249–59. doi:10.1016/j.immuni.2011.08.008

8. Ahmadi F, Junghus F, Ashworth C, et al. cDC1-derived IL-27 regulates small intestinal CD4+ T cell homeostasis in mice. J Exp Med. Mar 6 2023;220(3)doi:10.1084/jem.20221090

9. Luda KM, Joeris T, Persson EK, et al. IRF8 Transcription-Factor-Dependent Classical Dendritic Cells Are Essential for Intestinal T Cell Homeostasis. Immunity. Apr 19 2016;44(4):860–74. doi:10.1016/j.immuni.2016.02.008

10. Deets KA, Nichols Doyle R, Rauch I, Vance RE. Inflammasome activation leads to cDC1-independent cross-priming of CD8 T cells by epithelial cell-derived antigen. Elife. Dec 23 2021;10 doi:10.7554/eLife.72082

11. Gao Y, Nish SA, Jiang R, et al. Control of T helper 2 responses by transcription factor IRF4-dependent dendritic cells. Immunity. Oct 17 2013;39(4):722–32. doi:10.1016/j.immuni.2013.08.028

12. Persson EK, Uronen-Hansson H, Semmrich M, et al. IRF4 transcription-factor-dependent CD103(+)CD11b(+) dendritic cells drive mucosal T helper 17 cell differentiation. Immunity. May 23 2013;38(5):958–69. doi:10.1016/j.immuni.2013.03.009

13. Schlitzer A, McGovern N, Teo P, et al. IRF4 transcription factor-dependent CD11b+ dendritic cells in human and mouse control mucosal IL-17 cytokine responses. Immunity. May 23 2013;38(5):970–83. doi:10.1016/j.immuni.2013.04.011

14. Akagbosu B, Tayyebi Z, Shibu G, et al. Novel antigen-presenting cell imparts T(reg)-dependent tolerance to gut microbiota. Nature. Oct 2022;610(7933):752–760. doi:10.1038/s41586-022-05309-5

15. Kotloff KL, Nataro JP, Blackwelder WC, et al. Burden and aetiology of diarrhoeal disease in infants and young children in developing countries (the Global Enteric Multicenter Study, GEMS): a prospective, case-control study. Lancet. Jul 20 2013;382(9888):209–22. doi:10.1016/S0140-6736(13)60844-2

16. Khalil IA, Troeger C, Rao PC, et al. Morbidity, mortality, and long-term consequences associated with d i arrhoea f rom Cryptosporidium infection in children younger than 5 years: a meta-analyses study. Lancet Glob Health. Jul 2018;6(7):e758–e768. doi:10.1016/S2214-109X(18)30283-3

17. Manabe YC, Clark DP, Moore RD, et al. Cryptosporidiosis in patients with AIDS: correlates of disease and survival. Clin Infect Dis. Sep 1998;27(3):536–42. doi:10.1086/514701

18. Gerber DA, Green M, Jaffe R, Greenberg D, Mazariegos G, Reyes J. Cryptosporidial infections after solid organ transplantation in children. Pediatr Transplant. Feb 2000;4(1):50–5. doi:10.1034/j.1399-3046.2000.00087.x

19. Mosier DA, Oberst RD. Cryptosporidiosis. A global challenge. Ann NY Acad Sci. 2000; 916: 102–11. doi: 10.1111/j.1749-6632.2000.tb05279.x

20. Chen XM, Keithly JS, Paya CV, LaRusso NF. Cryptosporidiosis. The New England journal of medicine. May 30 2002;346(22):1723–31. doi:10.1056/NEJMra013170

21. O’Connor R M, Shaffie R, Kang G, Ward HD. Cryptosporidiosis in patients with HIV/AIDS. AIDS. Mar 13 2011;25(5):549–60. doi:10.1097/QAD.0b013e3283437e88

22. Okhuysen PC, Chappell CL. Cryptosporidium virulence determinants--are we there yet? Int J Parasitol. May 2002;32(5):517–25. doi:10.1016/s0020-7519(01)00356-3

23. Bouzid M, Hunter PR, Chalmers RM, Tyler KM. Cryptosporidium pathogenicity and virulence. Clin Microbiol Rev. Jan 2013;26(1):115–34. doi:10.1128/CMR.00076-12

24. Guerin A, Striepen B. The Biology of the Intestinal Intracellular Parasite Cryptosporidium. Cell host & microbe. Oct 7 2020;28(4):509–515. doi:10.1016/j.chom.2020.09.007

25. McDonald V, Bancroft GJ. Mechanisms of innate and acquired resistance to Cryptosporidium parvum infection in SCID mice. Parasite Immunol. Jun 1994;16(6):315–20. doi:10.1111/j.1365-3024.1994.tb00354.x

26. Pollok RC, Farthing MJ, Bajaj-Elliott M, Sanderson IR, McDonald V. Interferon gamma induces enterocyte resistance against infection by the intracellular pathogen Cryptosporidium parvum. Gastroenterology. Jan 2001;120(1):99–107. doi:10.1053/gast.2001.20907

27. Gomez Morales MA, Mele R, Ludovisi A, et al. Cryptosporidium parvum-specific CD4 Th1 cells from sensitized donors responding to both fractionated and recombinant antigenic proteins. Infection and immunity. Mar 2004; 72 (3) : 1306–10. doi: 10.1128/iai.72.3.1306-1310.2004

28. Ehigiator HN, McNair N, Mead JR. Cryptosporidium parvum: the contribution of Th1-inducing pathways to the resolution of infection in mice. Exp Parasitol. Feb 2007;115(2):107–13. doi:10.1016/j.exppara.2006.07.001

29. Borad A, Ward H. Human immune responses in cryptosporidiosis. Future microbiology. Mar 2010;5(3):507–19. doi:10.2217/fmb.09.128

30. McNair NN, Mead JR. CD4(+) effector and memory cell populations protect against Cryptosporidium parvum infection. Microbes Infect. Jul-Aug 2013;15(8-9):599–606. doi:10.1016/j.micinf.2013.04.009

31. Sateriale A, Slapeta J, Baptista R, et al. A Genetically Tractable, Natural Mouse Model of Cryptosporidiosis Offers Insights into Host Protective Immunity. Cell host & microbe. Jul 10 2019;26(1):135–146 e5. doi:10.1016/j.chom.2019.05.006

32. Gullicksrud JA, Sateriale A, Engiles JB, et al. Enterocyte-innate lymphoid cell crosstalk drives early IFN-gamma-mediated control of Cryptosporidium. Mucosal Immunol. Nov 8 2021;doi:10.1038/s41385-021-00468-6

33. Ungar BL, Kao TC, Burris JA, Finkelman FD. Cryptosporidium infection in an adult mouse model. Independent roles for IFN-gamma and CD4+ T lymphocytes in protective immunity. J Immunol. Aug 1 1991;147(3):1014–22.

34. Cohn IS, Henrickson SE, Striepen B, Hunter CA. Immunity to Cryptosporidium: Lessons from Acquired and Primary Immunodeficiencies. J Immunol. Dec 15 2022;209(12):2261–2268. doi:10.4049/jimmunol.2200512

35. Lantier L, Lacroix-Lamande S, Potiron L, et al. Intestinal CD103+ dendritic cells are key players in the innate immune control of Cryptosporidium parvum infection in neonatal mice. PLoS Pathog. 2013;9(12):e1003801. doi:10.1371/journal.ppat.1003801

36. Potiron L, Lacroix-Lamande S, Marquis M, et al. Batf3-Dependent Intestinal Dendritic Cells Play a Critical Role in the Control of Cryptosporidium parvum Infection. The Journal of infectious diseases. Feb 23 2019;219(6):925–935. doi:10.1093/infdis/jiy528

37. Russler-Germain EV, Jung J, Miller AT, et al. Commensal Cryptosporidium colonization elicits a cDC1-dependent Th1 response that promotes intestinal homeostasis and limits other infections. Immunity. Oct 21 2021; doi: 10.1016/j.immuni.2021.10.002

38. Wang HC, Zhou Q, Dragoo J, Klein JR. Most murine CD8+ intestinal intraepithelial lymphocytes are partially but not fully activated T cells. J Immunol. Nov 1 2002;169(9):4717–22. doi:10.4049/jimmunol.169.9.4717

39. Klein JR. T-cell activation in the curious world of the intestinal intraepithelial lymphocyte. Immunol Res. 2004;30(3):327–37. doi:10.1385/IR:30:3:327

40. Montufar-Solis D, Garza T, Klein JR. T-cell activation in the intestinal mucosa. Immunol Rev. Feb 2007;215:189–201. doi:10.1111/j.1600-065X.2006.00471.x

41. Vandereyken M, James OJ, Swamy M. Mechanisms of activation of innate-like intraepithelial T lymphocytes. Mucosal Immunol. Sep 2020;13(5):721–731. doi:10.1038/s41385-020-0294-6

42. Dumaine JE, Sateriale A, Gibson AR, et al. The enteric pathogen Cryptosporidium parvum exports proteins into the cytosol of the infected host cell. Elife. Dec 6 2021;10 doi:10.7554/eLife.70451

43. Haskins BE, Gullicksrud JA, Wallbank BA, et al. Dendritic cell-mediated responses to secreted Cryptosporidium effectors are required for parasite-specific CD8 (+) T cell responses. bioRxiv. Aug 18 2023;doi:10.1101/2023.08.16.553566

44. Moon JJ, Chu HH, Pepper M, et al. Naive CD4(+) T cell frequency varies for different epitopes and predicts repertoire diversity and response magnitude. Immunity. Aug 2007;27(2):203–13. doi:10.1016/j.immuni.2007.07.007

45. Oxenius A, Bachmann MF, Zinkernagel RM, Hengartner H. Virus-specific MHC-class II-restricted TCR-transgenic mice: effects on humoral and cellular immune responses after viral infection. Eur J Immunol. Jan 1998; 28 (1): 390–400. doi: 10.1002/(SICI)1521-4141(199801)28:01<390::AID-IMMU390>3.0.CO;2-O

46. Harrington LE, Janowski KM, Oliver JR, Zajac AJ, Weaver CT. Memory CD4 T cells emerge from effector T-cell progenitors. Nature. Mar 20 2008;452(7185):356–60. doi:10.1038/nature06672

47. Kato N, Comer E, Sakata-Kato T, et al. Diversity-oriented synthesis yields novel multistage antimalarial inhibitors. Nature. Oct 20 2016;538(7625):344–349. doi:10.1038/nature19804

48. Vinayak S, Jumani RS, Miller P, et al. Bicyclic azetidines kill the diarrheal pathogen Cryptosporidium in mice by inhibiting parasite phenylalanyl-tRNA synthetase. Sci Transl Med. Sep 30 2020;12(563)doi:10.1126/scitranslmed.aba8412

49. Shaw S, Cohn IS, Baptista RP, et al. Genetic crosses within and between species of Cryptosporidium. bioRxiv. Aug 4 2023;doi:10.1101/2023.08.04.551960

50. Everts B, Tussiwand R, Dreesen L, et al. Migratory CD103+ dendritic cells suppress helminth-driven type 2 immunity through constitutive expression of IL-12. J Exp Med. Jan 11 2016;213(1):35–51. doi:10.1084/jem.20150235

51. Moran AE, Holzapfel KL, Xing Y, et al. T cell receptor signal strength in Treg and iNKT cell development demonstrated by a novel fluorescent reporter mouse. J Exp Med. Jun 6 2011;208(6):1279–89. doi:10.1084/jem.20110308

52. Parsa R, London M, Rezende de Castro TB, et al. Newly recruited intraepithelial Ly6A(+)CCR9(+)CD4(+) T cells protect against enteric viral infection. Immunity. Jul 12 2022;55(7):1234–1249 e6. doi:10.1016/j.immuni.2022.05.001

53. Ungar BL, Burris JA, Quinn CA, Finkelman FD. New mouse models for chronic Cryptosporidium infection in immunodeficient hosts. Infection and immunity. Apr 1990;58(4):961–9. doi:10.1128/iai.58.4.961-969.1990

54. Pardy RD, Wallbank BA, Striepen B, Hunter CA. Immunity to Cryptosporidium: insights into principles of enteric responses to infection. Nat Rev Immunol. Sep 11 2023;doi:10.1038/s41577-023-00932-3

55. Vinayak S, Pawlowic MC, Sateriale A, et al. Genetic modification of the diarrhoeal pathogen Cryptosporidium parvum. Nature. Jul 23 2015;523(7561):477–80. doi:10.1038/nature14651

56. Murphy KM, Reiner SL. The lineage decisions of helper T cells. Nat Rev Immunol. Dec 2002;2(12):933–44. doi:10.1038/nri954

57. Trinchieri G. Interleukin-12 and the regulation of innate resistance and adaptive immunity. Nat Rev Immunol. Feb 2003;3(2):133–46. doi:10.1038/nri1001

58. Gazzinelli RT, Wysocka M, Hayashi S, et al. Parasite-induced IL-12 stimulates early IFN-gamma synthesis and resistance during acute infection with Toxoplasma gondii. J Immunol. Sep 15 1994;153(6):2533–43.

59. Tripp CS, Kanagawa O, Unanue ER. Secondary response to Listeria infection requires IFN-gamma but is partially independent of IL-12. J Immunol. Oct 1 1995;155(7):3427–32.

60. Park AY, Hondowicz BD, Scott P. IL-12 is required to maintain a Th1 response during Leishmania major infection. J Immunol. Jul 15 2000;165(2):896–902. doi:10.4049/jimmunol.165.2.896

61. Feng CG, Jankovic D, Kullberg M, et al. Maintenance of pulmonary Th1 effector function in chronic tuberculosis requires persistent IL-12 production. J Immunol. Apr 1 2005;174(7):4185–92. doi:10.4049/jimmunol.174.7.4185

62. Christian DA, Adams TA, 2nd, Shallberg LA, et al. cDC1 coordinate innate and adaptive responses in the omentum required for T cell priming and memory. Sci Immunol. Sep 30 2022;7(75):eabq7432. doi:10.1126/sciimmunol.abq7432

63. Liesenfeld O, Kosek J, Remington JS, Suzuki Y. Association of CD4+ T cell-dependent, interferon-gamma-mediated necrosis of the small intestine with genetic susceptibility of mice to peroral infection with Toxoplasma gondii. J Exp Med. Aug 1 1996;184(2):597–607. doi:10.1084/jem.184.2.597

64. Ruterbusch MJ, Hondowicz BD, Takehara KK, Pruner KB, Griffith TS, Pepper M. Allergen exposure functionally alters influenza-specific CD4+ Th1 memory cells in the lung. J Exp Med. Nov 6 2023;220(11)doi:10.1084/jem.20230112

65. Devarajan P, Vong AM, Castonguay CH, et al. Cytotoxic CD4 development requires CD4 effectors to concurrently recognize local antigen and encounter type I IFN-induced IL-15. Cell Rep. Sep 29 2023;42(10):113182. doi:10.1016/j.celrep.2023.113182

66. Hildner K, Edelson BT, Purtha WE, et al. Batf3 deficiency reveals a critical role for CD8alpha+ dendritic cells in cytotoxic T cell immunity. Science. Nov 14 2008;322(5904):1097–100. doi:10.1126/science.1164206

67. Durai V, Bagadia P, Granja JM, et al. Cryptic activation of an Irf8 enhancer governs cDC1 fate specification. Nat Immunol. Sep 2019;20(9):1161–1173. doi:10.1038/s41590-019-0450-x

68. Hsieh CS, Macatonia SE, Tripp CS, Wolf SF, O’Garra A, Murphy KM. Development of TH1 CD4+ T cells through IL-12 produced by Listeria-induced macrophages. Science. Apr 23 1993;260(5107):547–9. doi:10.1126/science.8097338

69. Cosyns M, Tsirkin S, Jones M, Flavell R, Kikutani H, Hayward AR. Requirement of CD40-CD40 ligand interaction for elimination of Cryptosporidium parvum from mice. Infection and immunity. Feb 1998;66(2):603–7. doi:10.1128/IAI.66.2.603-607.1998

70. Wu R, Murphy KM. DCs at the center of help: Origins and evolution of the three-cell-type hypothesis. J Exp Med. Jul 4 2022;219(7)doi:10.1084/jem.20211519

71. Maria C.C. Canesso TBRC, Sandra Nakandakari-Higa, Ainsley Lockhart, Daria Esterházy, Bernardo S. Reis, Gabriel D. Victora, Daniel Mucida. Identification of dendritic cell-T cell interactions driving immune responses to food. bioRxiv. 2023;

72. Bilate AM, London M, Castro TBR, et al. T Cell Receptor Is Required for Differentiation, but Not Maintenance, of Intestinal CD4(+) Intraepithelial L ymphocytes. Immunity. Nov 17 2020;53(5):1001–1014 e20. doi:10.1016/j.immuni.2020.09.003

73. Ryan D. Pardy KAW, Bethan A. Wallbank, Jessica H. Byerly, Keenan M. Ian S. Cohn, Breanne E. Haskins, Justin L. Roncaioli, Eleanor J. Smith, Gracyn Y. Buenconsejo, Boris Striepen and Christopher A. Hunter. Delayed intestinal epithelial cell responsiveness to IFN-γ mediates the timing of parasite control during Cryptosporidium infection. bioRxiv. 2013;

74. Choudhry N, Petry F, van Rooijen N, McDonald V. A protective role for interleukin 18 in interferon gamma-mediated innate immunity to Cryptosporidium parvum that is independent of natural killer cells. The Journal of infectious diseases. Jul 1 2012;206(1):117–24. doi:10.1093/infdis/jis300

75. Sateriale A, Gullicksrud JA, Engiles JB, et al. The intestinal parasite Cryptosporidium is controlled by an enterocyte intrinsic inflammasome that depends on NLRP6. Proceedings of the National Academy of Sciences of the United States of America. Jan 12 2021;118(2)doi:10.1073/pnas.2007807118

76. Zhao GH, Fang YQ, Ryan U, et al. Dynamics of Th17 associating cytokines in Cryptosporidium parvum-infected mice. Parasitol Res. Feb 2016;115(2):879–87. doi:10.1007/s00436-015-4831-2

77. Drinkall E, Wass MJ, Coffey TJ, Flynn RJ. A rapid IL-17 response to Cryptosporidium parvum in the bovine intestine. Vet Immunol Immunopathol. Sep 2017; 191:1–4. doi: 10.1016/j.vetimm.2017.07.009

78. Kiner E, Willie E, Vijaykumar B, et al. Gut CD4(+) T cell phenotypes are a continuum molded by microbes, not by TH archetypes. Nat Immunol. Jan 18 2021;doi:10.1038/s41590-020-00836-7

79. Tandel J, English ED, Sateriale A, et al. Life cycle progression and sexual development of the apicomplexan parasite Cryptosporidium parvum. Nat Microbiol. Dec 2019;4(12):2226–2236. doi:10.1038/s41564-019-0539-x

80. Manjunatha UH, Vinayak S, Zambriski JA, et al. A Cryptosporidium PI(4)K inhibitor is a drug candidate for cryptosporidiosis. Nature. Jun 15 2017;546(7658):376–380. doi:10.1038/nature22337

81. Hirota K, Turner JE, Villa M, et al. Plasticity of Th17 cells in Peyer’s patches is responsible for the induction of T cell-dependent IgA responses. Nat Immunol. Apr 2013;14(4):372–9. doi:10.1038/ni.2552

